# Mutant screen for reproduction unveils depression-associated Piccolo’s control over reproductive behavior

**DOI:** 10.1101/405985

**Authors:** Gerardo A. Medrano, Manvendra Singh, Erik J. Plautz, Levi B. Good, Karen M. Chapman, Jaideep Chaudhary, Priscilla Jaichander, Heather M. Powell, Ashutosh Pudasaini, John M. Shelton, James A. Richardson, Xian-Jin Xie, Zoltán Ivics, Christine Braun, Frauke Ackermann, Craig C. Garner, Zsuzsanna Izsvák, F. Kent Hamra

## Abstract

Successful sexual reproduction involves complex, genetically encoded interplay between animal physiology and behavior. The rat provides a highly fecund mammalian model for studying how the brain impacts reproduction. Here, we report a forward genetics screen in rats to identify genes that affect reproduction. A panel of 18 distinct rat strains harboring *Sleeping Beauty* gene trap mutations were analyzed for the ability to reproduce. As expected, our mutant screen identified genes where reproductive failure was connected to gametogenesis (*Btrc, Pan3, Spaca6, Ube2k*) and embryogenesis (*Alk3, Exoc6b, Slc1a3, Tmx4, Zmynd8*). In addition, we identified *Atg13* (longevity) and *Pclo* (neuronal disorders), previously not associated with an inability to conceive. Neurologically, Pclo is known to regulate the size of presynaptic vesicle pools. Here, dominant traits in *Pclo* mutant rats caused epileptiform activity and affected genes supporting GABAergic synaptic transmission (*Gabra6*, *Gabrg3*). Recessive traits in *Pclo* mutant rats transmitted altered reproductive behavior, as homozygous *Pclo* mutant rats produced gametes but neither sex would mate with wildtype rats. *Pclo* mutant rat behavior was linked to endophenotypes signifying compromised brain-gonad crosstalk via disturbed GnRH signaling and allelic markers for major depressive disorder in humans (*Grm5, Htr2a, Sorcs3, Negr1, Drd2*). Thus, by rat genetics, we identified Pclo as a candidate presynaptic factor required for reproduction.

**Author Summary:** Piccolo gene mutations have previously been identified in human cohorts diagnosed with behavioral syndromes that impact one’s emotions, including depression and bipolar disorder. Although studies in human populations implicate changes to *Piccolo’s* DNA sequence to enhanced susceptibility for behavioral disorders, studies in mouse models have yet to link *Piccolo* mutations to altered behavior. Here, by a novel genetics approach, we report *Piccolo* mutation-dependent effects on reproductive behavior in rats, a finding that may turn out to be relevant to the behavioral effects that are associated with human *Piccolo* gene mutations. Thus, research aimed at understanding how *Piccolo* functions to regulate reproduction in rats could prove pivotal in our ability to understand neurological mechanisms that influence human emotions.

## Introduction

While a failure to reproduce sexually is often connected to physiological or developmental problems of the gonad, gamete or embryo, it is also commonly accepted that problems with sexual reproduction can be linked to physical fitness [1] or behavioral abnormalities [2]. Indeed, inborn social behaviors related to sex, defense and maternal care are elicited by sensory input that is processed by the central nervous system to promote successful reproduction [3].

Rats are highly fecund mammals and display robust appetitive and consummatory reproductive behavior [4, 5]. In rats, sensory input to the limbic system that drives reproduction is mediated in large part via the olfactory system [olfactory epithelia > olfactory nuclei > main and/or accessory olfactory bulb > medial amygdala > bed nucleus of stria terminalis > medial pre-optic hypothalamic nucleus and ventromedial hypothalamus] [3]. Pheromones that signal mating bind to chemosensory olfactory receptors in the olfactory epithelium to elicit pre-copulatory social behaviors such as partner investigation, grooming and courtship [3, 6, 7]. Pre-copulatory sensory signals further culminate in copulatory and post-copulatory behavior that enable fertilization [3, 6, 7]. Notably, the rat’s olfactory epithelium is uniquely endowed with ~1,400 genes encoding olfactory receptors [8] and has long provided an experimental system to study mechanisms by which sensory input stimulates social behavior responses that affect reproduction [3, 6, 7].

From the hundreds of genes essential for neuroendocrine/gonadal control over gametogenesis and fertilization [9], neurotransmission genes that govern sensory neuron-stimulated social behavior mediate the primary signals that initiate reproduction [3, 6, 7]. Social responses such as pleasure, attraction, fear, aggression and avoidance that affect reproduction are processed by the limbic system to modulate motivational responses [2, 10]. Innate reproductive behaviors are driven by afferent sensory neurons that innervate the limbic system in mammals and are driven by sex and sex hormones (estrogen and testosterone) [6, 7]. Abnormalities in the cortico-limbic networks that integrate survival-driven reproductive behavior with emotional awareness and memory play crucial roles in the etiology of human “affective disorders”, including depression, bipolar disorder, autism, anxiety and addiction, and represent neurological health conditions [11-13].

In this study, we aimed to identify novel genes required for reproduction. Our intention was to reach out from the circle of obvious candidates and uncover novel layers of reproductive biology, remaining as open as possible to finding the unexpected. Therefore, instead of taking a targeted approach, we chose an unbiased, forward mutagenesis strategy to identify new genes that impact reproduction using the rat model.

*Sleeping Beauty* transposon genomic insertions occur randomly, with ~35% frequency of landing in a gene transcribed by RNA polymerase II [14-16]. Gene trapping by random, *Sleeping Beauty-mediated* insertion of a selectable marker (e.g. *β-geo*) into RNA Polymerase II transcription units provides a powerful technology for introducing disruptive mutations into genes on a genome-wide scale. The ability to select for recombinant spermatogonial stem cell libraries harboring *Sleeping Beauty* transposon gene trap (gt) insertions has enabled large-scale production of novel mutant rat strains for analyses in forward genetic assays [17].

In the current study, a panel of *Sleeping Beauty* mutant rat strains derived from a spermatogonial gene trap library were tested for impaired reproduction phenotypes. In addition to genes required for gamete and embryo development, our mutant screen unveiled new genetic connections between reproduction, fitness and social behavior. Among the reproduction genes, we identified *Atg13,* which has generally been connected to longevity in species ranging from yeast to plants and humans. We also identified phenotypes in *Pclo*-deficient (*Pclo*^gt/gt^) rats that hold potential to model humans diagnosed with neurological disorders [18-20].

## Results

### A set of mutations affects reproduction

To identify genes essential for reproduction, we conducted a forward genetic screen using rats derived from a spermatogonial library of *Sleeping Beauty* insertional mutations [17] (**Fig 1**). Each mutant rat strain tested inherited a copy of *Sleeping Beauty* inserted within a distinct protein coding gene (n=17 gene trap insertions + 1 untrapped gene insertion) (**S1 Fig** and **S1 Table**). In total, 12 of 18 mutant genes analyzed proved to be essential for reproduction (**Fig 2A** and **S2 Table**). Inability to reproduce was linked to a variety of phenotypes that included gametogenesis defects (*Btrc*^gt/gt^, *Ube2k*^gt/gt^, *Pan3*^gt/gt^, *Spaca6*^gt/gt^), embryonic lethality (*Alk3*^gt/gt^, *Exoc6b*^gt/gt^, *Slc1a3*^gt/gt^, *Tmx4*^gt/gt^, *Zmynd8*^gt/gt^), end-stage organ failure (*Atg13*^gt/gt^) and impaired behavior (*Pclo*^gt/gt^, *Dlg1*^wt/gt^) (**S3 Table and S4 Table**).

**Figure 1.**
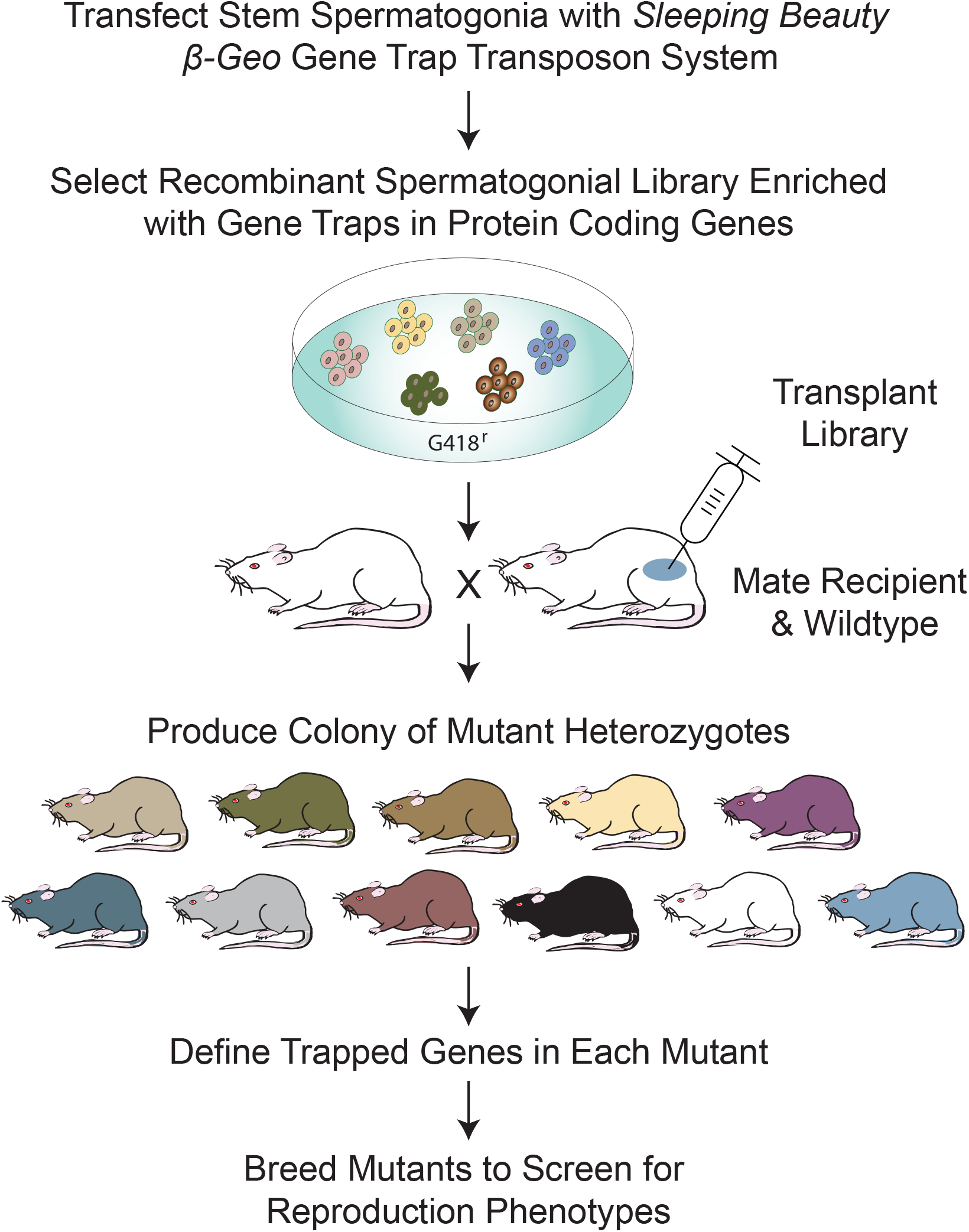
Sperm Stem Cell Based Forward Screen for Rat Reproduction Genes. Recombinant rat spermatogonial stem cell libraries are produced by *Sleeping Beauty* transposon genomic insertion. Spermatogonial libraries of randomly inserted *Sleeping Beauty* gene trap mutations are used to produce colonies of mutant rats. Novel *Sleeping Beauty* mutant rat strains are crossed to identify genes that impact reproduction. In the current study, eleven homozygous mutant rat strains generated were viable following birth (~70%), 6 were embryonic lethal (~28%) and 1 was scored as sub-viable postnatally (~6%) (n=18 mutant rat strains analyzed for ability to reproduce). Similar relative percentages were reported in mice by the European Conditional Mouse Mutagenesis Program (EUCOMM) and the Knockout Mouse Project (KOMP) [63].

**Figure 2.**
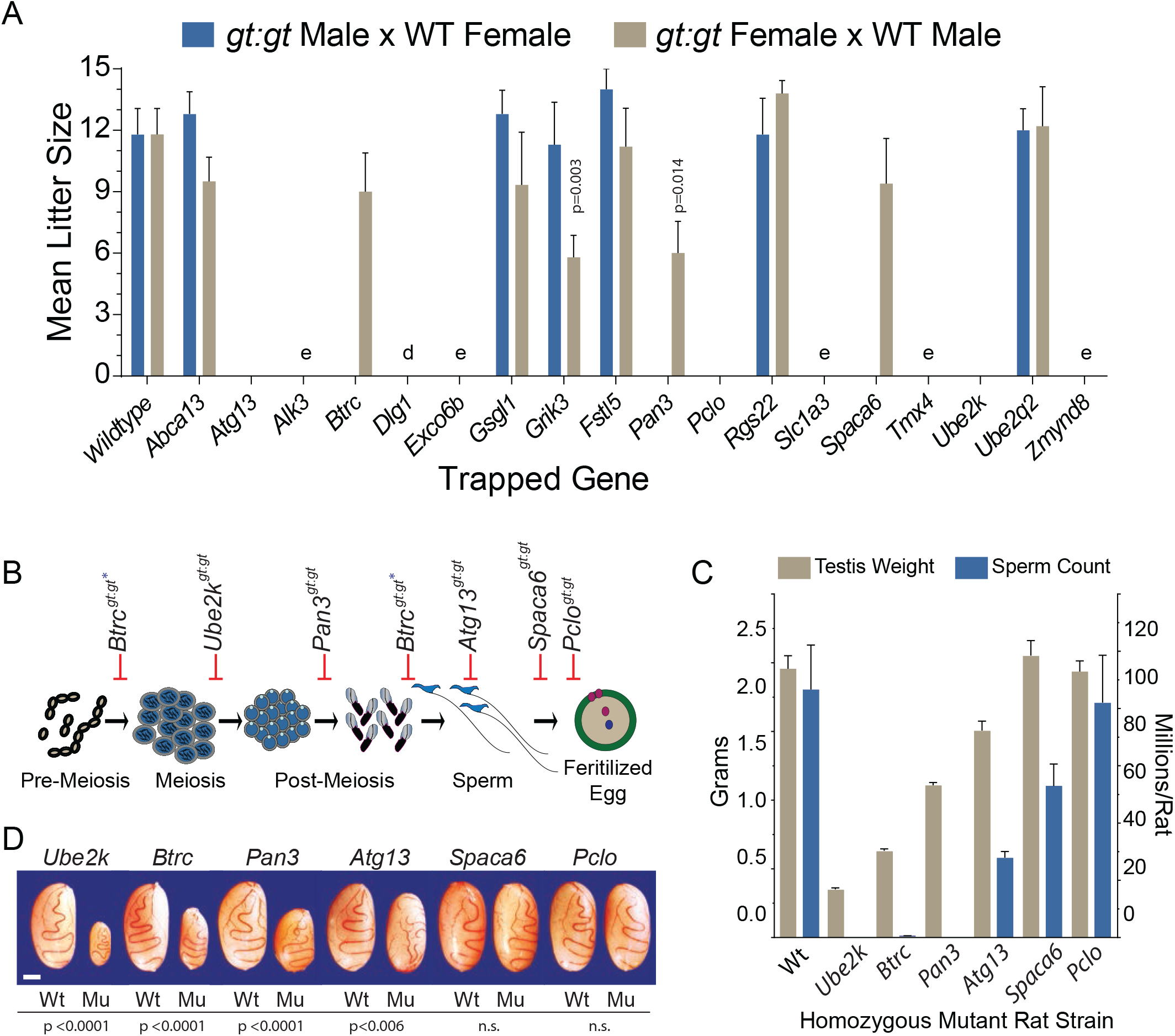
Gene Mutations that Cause Infertility in Rats. (**A**) Mean litter size produced by crossing female and male homozygous *Sleeping Beauty* mutant rats *(gt:gt*) with wildtype breeders (WT). e = embryonic lethal; d = dominant effect that disrupted F1 female mutant fecundity. (**B**) Developmental steps during sperm maturation or fertilization disrupted by respective homozygous gene trap mutations (*gt:gt*) in rats. *Note: *Btrc*^gt/gt^ rats displayed pre-meiotic (~85% tubules) and post-meiotic (~15% tubules) spermatogenic arrest based on co-labeling with nuclear markers (γH2AX and Hoechst 43332 dye). (**C**) Mean testis weight (tan bars; left y-axis) and epididymal sperm counts (blue bars; right y-axis) from respective homozygous mutant rat strains (±SEM, n=4-6 rats/strain). Measurements taken between postnatal days 120-180. Caudal epididymal spermatozoa from *Spaca6*^gt/gt^ (n=6) and *Pclo*^gt/gt^ (n=4) rats displayed similar basal activity compared to wildtype. (**D**) Testes from wildtype (Wt) and homozygous gene trap mutants (Mu). Scale bar, 5 mm

### Mutations that disrupt distinct steps in rat spermatogenesis

Homozygous gene trap mutations in *Btrc, Ube2k* and *Pan3* blocked spermatogenesis at pre-meiotic, meiotic and post-meiotic steps, respectively (**Fig 2B, S2A and S2B**). Only residual numbers of malformed spermatozoa were detected in *Btrc*^gt/gt^ males, and no epididymal spermatozoa were observed in *Ube2k*^gt/gt^ or *Pan3*^gt/gt^ males (**Fig 2C** and **S3 Table**). In corresponding *Ube2k*^gt/gt^, *Btrc*^gt/gt^ and *Pan3*^gt/gt^ genotypes, spermatogenic arrest was reflected by reduced testis size (**Fig 2D** and **S3 Table**). *Btrc, Ube2k* and *Pan3* provide a novel set of reproduction models, where the infertility is the result of developmental problems of the gonad.

### A group of mutant rats develop gametes, but do not reproduce

Rats with homozygous mutations in the *Spaca6, Atg13* and *Pclo* genes produced both eggs and sperm (**Fig 2C, S2A and S2C**). However, neither sex of *Atg13*^gt/gt^ and *Pclo*^gt/gt^ rats were able to reproduce, as was the case with *Spaca6*^gt/gt^ males (**Fig 2A** and **S2 Table**). *Spaca6*^gt/gt^ females produced relatively normal sized litters when paired with wt males (**Fig 2A** and **S2 Table**). Mating behavior appeared normal in *Spaca6*^gt/gt^ males when compared to wt males as supported by the presence of spermatozoa in vaginal swabs (n=4 breeder pairs). *Spaca6*^gt/gt^ males had normal size testes (**Fig 2D**). While *Spaca6*^gt/gt^ epididymides had slightly reduced numbers of spermatozoa (**Fig 2C**), their moderate deviation in sperm counts could not explain the observed infertility phenotype. The reproductive failure of *Spaca6*^gt/gt^ males might be more likely associated with the recently suggested role of this immunglobulin-like protein in regulating sperm-egg membrane fusion [21].

### Reproduction defects in *Atg13* mutants correlate with reduced longevity

Whereas *Autophagy related 13* (*Atg13*) is required for autophagic flux and reaching an optimal lifespan in plants and animals (**S4 Table**)[22-24], the role of *Atg13* in additional reproduction-related traits is unknown. The insertional mutation resulted in a truncated form of *Atg13* predicted to lack exon 16 (*Atg13*^Δe16^) (**Fig 3A**). *Atg* exon 16 encodes the 25 carboxyl-terminal amino acids in Atg13 (**Fig 3A**). Expression of *Atg13^Δe16^* generated a protein that resembled wt ATG13: it was abundant in testes, with lower levels in other tissues (**Fig 3A inset**). All male *Atg13*^gt/gt^ mutants were characterized by reduced testis size and epididymal sperm counts compared to wt (**Fig 2C, D**) but had relatively high testis-to-body weight and epididymis-to-body weight ratios (**S3 Table**). *Atg13*^gt/gt^ cauda epididymal spermatozoa flagella were immotile and displayed more detached heads and tails than WT (n=4/genotype).

**Figure 3.**
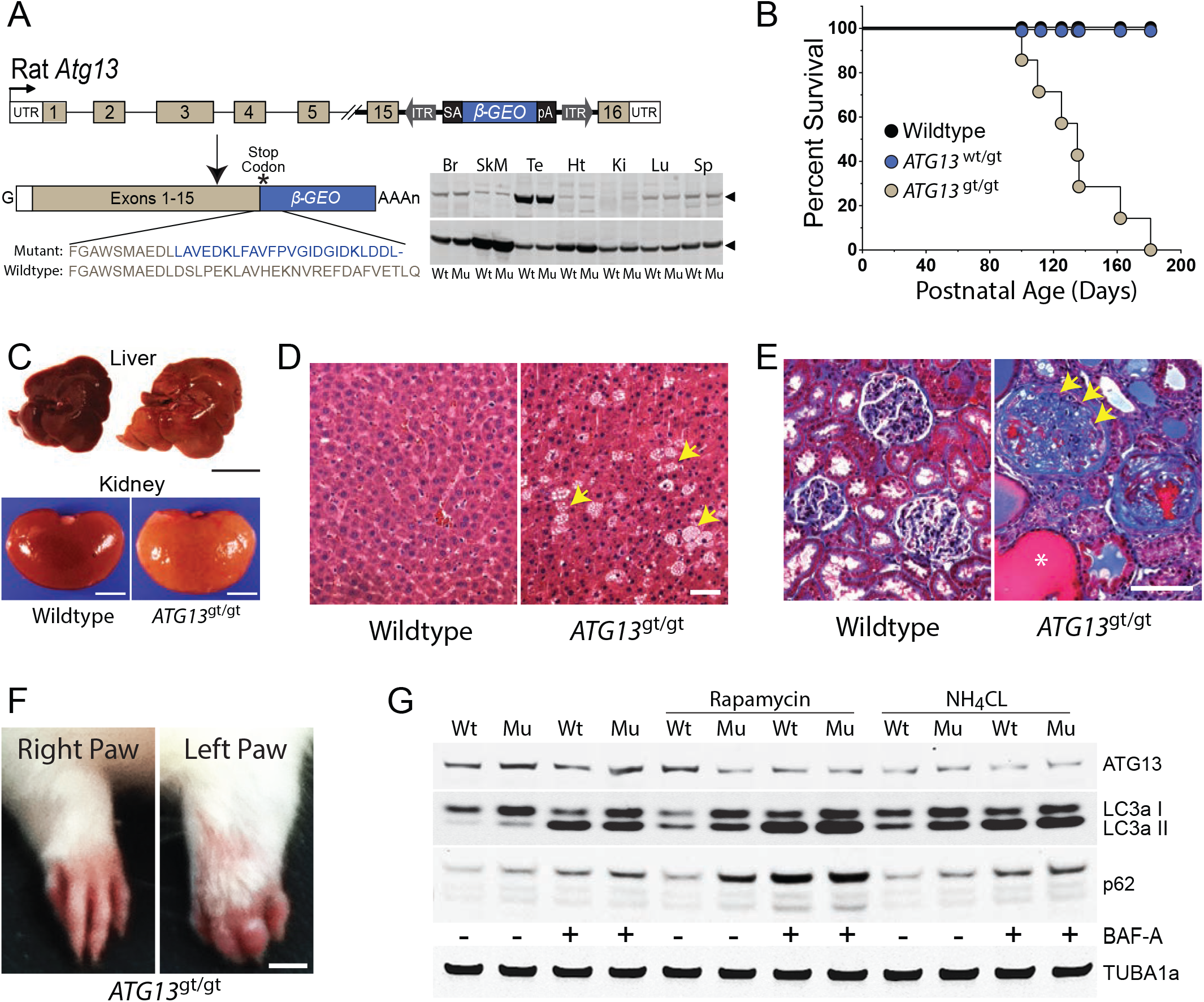
Pathology linked to the COOH-terminal 25 amino acids in rat Autophagy Related 13. (**A**) Diagram of *Sleeping Beauty β-Geo* gene trap in rat *Atg13* intron 15. The gene trap splices to *Atg13* exon 15 and out of frame with the *β-Geo* reporter that replaces *Atg13* exon 16 (*Atg13*^Δe16^). *Atg13* exon 16 encodes the C-terminal 25aa of wildtype Atg13 and is predicted to be replaced by a 24aa epitope (blue font) derived from the gene trap construct, thereby, generating a similar size mutant protein. *<u>Inset</u>*: (top panel) western blot probing ATG13 in tissues from wildtype (WT) and homozygous mutant *Atg13*^gt:gt^ (Mu) rat littermates; (bottom panel) same blot probed for Gapdh. Arrowheads point to WT and Mu rat proteins with molecular size of ATG13 (~65 kDa) and Gadph (~37 kDa). Br, brain; SkM, skeletal muscle; Te, testis; Ht, heart; Ki, kidney; Lu, lung; Sp, spleen (**B**) Kaplan-Meier estimator of postnatal survival for *Atg13*^wt:wt^ (wildtype), *Atg13*^wt:gt^ (heterozygous) and *Atg13*^gt:gt^ (homozygous) mutant rats. (**C**) Liver (top) and Kidney (bottom) from wildtype and homozygous mutant (*Atg13*^gt:gt^) littermates. Liver scale bar, 2 cm; Kidney scale bar 5 mm. (**D**) Hematoxylin and Eosin stained liver sections in *Atg13*^wt:wt^ and *Atg13*^gt:gt^ rats. Note, fatty liver in *Atg13*^gt:gt^ rats (arrows). Liver sections from littermates, postnatal day 110. Scale bar 50 μm. (**E**) Trichrome stained sections illustrating dramatic sclerosis of the glomerular tuft and fibrosis in Bowman’s capsule of an *Atg13*^gt:gt^ rat. Note proliferating epithelial cells lining Bowman’s capsule (arrows). An adjacent tubule is dilated and filled with protein rich filtrate (asterisks). Kidney sections from wildtype and *Atg13*^gt:gt^ littermates, postnatal D110. Scale bar 100 μm. (**F**) Forearms of *Atg13*^gt:gt^ phenotype in one strain. Note swelling of left arm/digits. Scale, 5 mm. (**G**) Relative expression of the autophagy marker proteins Atg13, LC3a I, LC3a II and p62 compared to TUBA1a (loading control) in embryonic fibroblasts derived from wildtype and mutant rats following treatment with or without combinations of rapamycin, ammonium chloride (NH_4_CL) and bafilomycin-A1 (BAF-A)

Beside the abnormal spermatozoa of males, all *Atg13*^gt/gt^ rats inherited pathologies associated with premature death at 3-5 months of age (**Fig 3B**). The livers and kidneys of *Atg13*^gt/gt^ rats were abnormal (**Fig 3C**), with the liver containing cells scattered throughout histological sections displaying small spherical vacuoles, consistent with an accumulation of triglycerides (**Fig 3D**). All the kidneys that were examined displayed marked glomerulonephritis and moderate tubule interstitial disease (**Fig 3E**). Homozygous *Atg13*^gt/gt^ animals (n=3) from one of three pedigrees also demonstrated edematous paws and digits in adult animals (**Fig 3F**).

Consistent with *Atg13*’s biological function, changes in the relative abundance of autophagy markers LC3a-I/II and p62 were found in *Atg13*^gt/gt^ embryonic fibroblasts (**Fig 3G**). Rapamycin treatment synergized with *Atg13*^gt/gt^ to increase LC3a-I/II and p62 relative abundance in fibroblasts, implicating Atg13’s COOH-terminal peptide in regulating mTorc-dependent autophagy signals (**Fig 3G**). The reproduction defects in both female and male *Atg13*^gt/gt^ rats correlated with adult-lethal pathologies, and in males, *Atg13*^gt/gt^ was further complicated with abnormal spermatozoa. Thus, the cause of infertility of *Atg13*^gt/gt^ rats is a cumulation of general fitness problems.

### Compromised neurotransmission in *Pclo* mutants

Despite their reproductive failure (**Fig 2A**), *Pclo*^gt/gt^ mutant rats did not display any obvious dysfunction during gametogenesis (**S2A** and **S2C Fig**). Numbers of epididymal spermatozoa from *Pclo*^gt/gt^ rats were relatively normal (**Fig 2C**). However, spermatozoa from *Pclo*^gt/gt^ rats were not found in vaginal swabs of WT females (6 of 6 breeder pairs) (**Fig 4A**, and spermatozoa from WT males were not detected in *Pclo*^gt/gt^ females (6 of 6 breeder pairs) (**Fig 4A**), suggesting that the mating did not actually occur.

**Figure 4.**
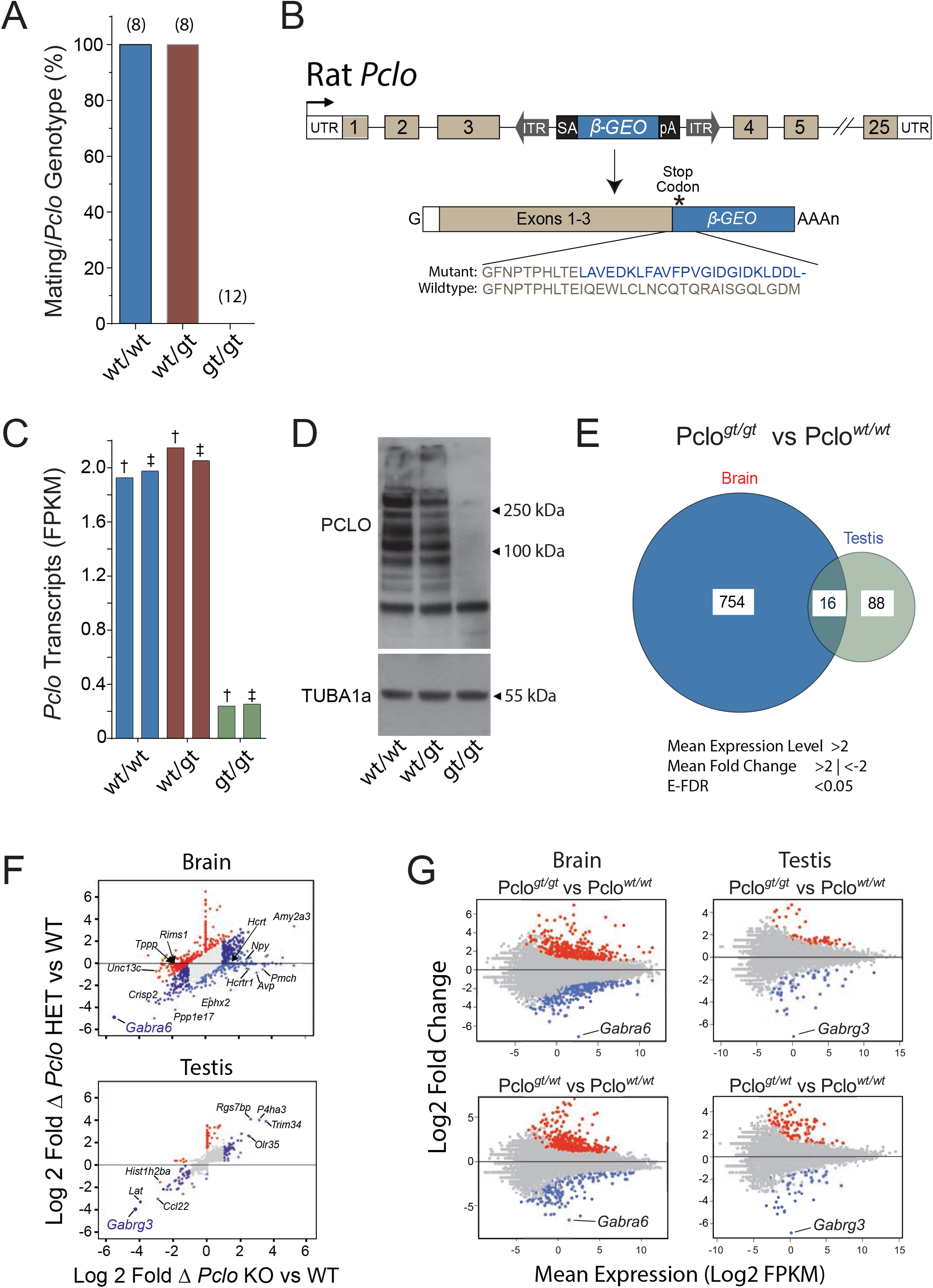
RNA profiling in mutant *Piccolo* rats. (**A**) *Pclo*^wt:wt^, *Pclo*^wt:gt^ and *Pclo*^gt:gt^ rat mating after pairing with wildtype breeders based on identification of spermatozoa in vaginal swabs. (n) = 8 to 12 total breeder pairs/genotype, or 4 to 6 breeder pairs/sex/genotype for *Pclo*^wt:gt^ and *Pclo*^gt:gt^ mutant strains. (**B**) Diagram of *Sleeping Beauty β-Geo* gene trap in rat *Pclo* intron 3. The gene trap splices to *Pclo* exon 3, out of frame with the *β-Geo* reporter. The gene trap is predicted to replace the C-terminal 3805aa or 4010aa encoded by exons 4-25 of respective wildtype *Pclo* isoforms with a 24aa construct-derived epitope (blue font) to generate *Pclo*^Δe4-25^. (**C**) Relative abundance (FPKM values) of *Pclo* transcript isoforms in *Pclo*^wt:wt^, *Pclo*^wt:gt^ and *Pclo*^gt:gt^ rat brains. ^†^NM_020098, encodes the full length 4880-amino acid isoform; ^‡^NM_001110797, encodes the full length 5041-amino acid isoform. (**D**) Western blot of Piccolo isoforms and TUBA1a in total brain lysates prepared from *Pclo*^wt:wt^, *Pclo*^wt:gt^ and *Pclo*^gt:gt^ rats. (**E**) Venn diagram shows the number of differentially expressed genes (DEGs) in the brain (754) and testis (88) or genes commonly expressed in both tissues (16) of *Pclo*^gt/gt^ rats compared to *Pclo*^wt/wt^ rats. (**F**) Relative abundance of DEGs in *Pclo*^wt/gt^ (HET) and *Pclo*^gt/gt^ (KO) rat brain (top) and testis (bottom) vs wildtype (WT). DEGs that changed more in HET or more in KO vs WT are shown in red and light blue, respectively (log2-fold change >1 or <-1; FDR < 0.05). Genes that changed comparably in abundance in both HET and KO but were differentially expressed relative to WT are shown in dark blue (log2-fold change >1 or <-1; FDR < 0.05). Note, that the *Pclo* mutation affected more changes in the brain vs testis transcriptome. (**G**) Fold change in relative brain (left) and testis (right) transcript abundance (Log2 FPKM values) in *Pclo*^wt/gt^ and *Pclo*^gt/gt^ rats vs *Pclo*^wt/wt^ rats. DEGs are shown in red (increased abundance) and blue (decreased abundance), respectively (log2-fold change >1 or <-1; FDR < 0.05). Note, the decreased *Gabra6* and *Gabra3* abundance in brain and testis, respectively, in both *Pclo*^wt/gt^ and *Pclo*^gt/gt^ rats.

*Pclo*^gt/gt^ rats harbor the *Sleeping Beauty* gene trap in *Pclo* intron 3, deleting exons 4-25 (*Pclo*^SBΔ4-25^ rats) (**Fig 4B**). *Pclo* encodes multiple protein isoforms (70-560kDa) that are primarily localized in the cytomatrix of pre-synaptic neurons and have been implicated to play a key role in synaptic transmission [25]. While Piccolo is expressed in various tissues, including testis, it is dominantly enriched in the brain, where it is relatively abundant in the cerebellum, pituitary gland, cortex, hypothalamus and nucleus accumbens (<u>GTEX Portal_PCLO</u>).

The unconsummated mating alongside with the dominant distribution of *Pclo* transcripts in the brain let us hypothesize that the reproductive failure in *Pclo*^gt/gt^ rats was associated by neurological abnormalities. Thus, to gain insights into Piccolo’s role in reproductive phenotypes, we used samples from both testes and brain tissues of *Pclo*^gt/gt^, *Pclo*^gt/wt^ and *Pclo*^wt/wt^ animals (~6 mo old), and subjected them to RNA sequencing (RNA-seq). As expected, *Pclo* transcripts were readily detected in the brain (**Fig 4C**), whereas the testicular expression of *Pclo* was low (< 0.1 FPKM) (**S5 Table**). In the brain, the gene trap insertion reduced *Pclo* transcript abundance by >8.5-fold in homozygous *Pclo*^gt/gt^ rats (**Fig 4C**), while no significant transcriptional changes of *Pclo* could be detected in heterozygous *Pclo*^wt/gt^ rats (**Fig 4C**). Similarly, at the protein level, Pclo was reduced by >95% in the brains of *Pclo*^gt/gt^ rats, but Pclo was not significantly affected in *Pclo*^wt/gt^ compared to *Pclo*^wt/wt^ littermates (**Fig 4D**). Thus, expression from a single *Pclo* allele appears to drive relatively normal levels of the gene product, and the phenotype that was observed appears to be connected to the allelic origin of Piccolo.

Our transcriptome analysis of *Pclo*^gt/gt^ and *Pclo*^wt/wt^ rats revealed a higher number of differentially expressed genes (DEGs) in the brain (754) than testis (88), while 16 genes were affected in both tissues (FPKM > 2 and log2 fold change |1| and E-FDR < 0.01) (**Fig 4E** and **S5 Table**). Inclusive to the 16 DEGs that were affected in both brain and testes, *Tspo, Ces1d, Folr1* and *Adh1* (**Fig 4E** and **S5 Table**) regulate steroid hormone/vitamin biosynthesis, signaling and transport in the blood stream [26-29]. Despite similar Piccolo RNA/protein abundance in *Pclo*^wt/wt^ and *Pclo*^wt/gt^ rat brains, 325 genes were differentially expressed (log2FC |1|) in the brains of heterozygotes compared to wildtype or homozygotes (**Fig 4F** and **S5 Table**), reflecting robust allelic effects.

Gene Ontology (GO) analyses of DEGs revealed the most significantly down-regulated processes in *Pclo*^gt/gt^ vs *Pclo*^wt/wt^ rats included *Synaptic Transmission* and *Neurogenesis* gene sets (p<0.000006; **S6 Table**), consistent with our hypothesis that lack of reproduction by *Pclo*^gt/gt^ rats was associated with a neurological defect. A prominent cluster of 80 downregulated *Synaptic Transmission* genes in the brain (**Fig 5A**) included the *gamma-aminobutyric acid (GABA) signaling pathway* (GO:0007214, p=0.0000009) (**Fig 5B** and **S6 Table**). The most significantly affected genes in both *Pclo*^gt/gt^ and *Pclo*^wt/gt^ rats were members of the GABA(A) receptor gene family, *Gabra6 (GABA(A) Receptor Subunit Alpha 6)* in the brain and *Gabrg3 (GABA(A) Receptor Subunit Gamma-3)* in the testis (**Fig 4G**). Notably, the expression of both, *Gabra6* in brain, and *Gabrg3* in testes, dropped to undetectable levels (FPKM < 0.01) in *Pclo*^wt/gt^ and *Pclo*^gt/wt^ rats, suggesting a dominant phenotype in *Pclo* ^gt/gt^ mutants that results in a close-to KO phenotype for each GABA(A) receptor subunit in respective tissues (**Fig 4G**). While *Gabra6* is dominantly expressed in the cerebellum of the brain (<u>GTEX Portal_Gabra6, FPKM >1</u>), *Gabrg3* is expressed in various regions of the brain, primarily in the hypothalamus, in the pituitary gland and has a moderate enrichment in the testis (<u>GTEX Portal_Gabrg3, FPKM>1</u>).

**Figure 5.**
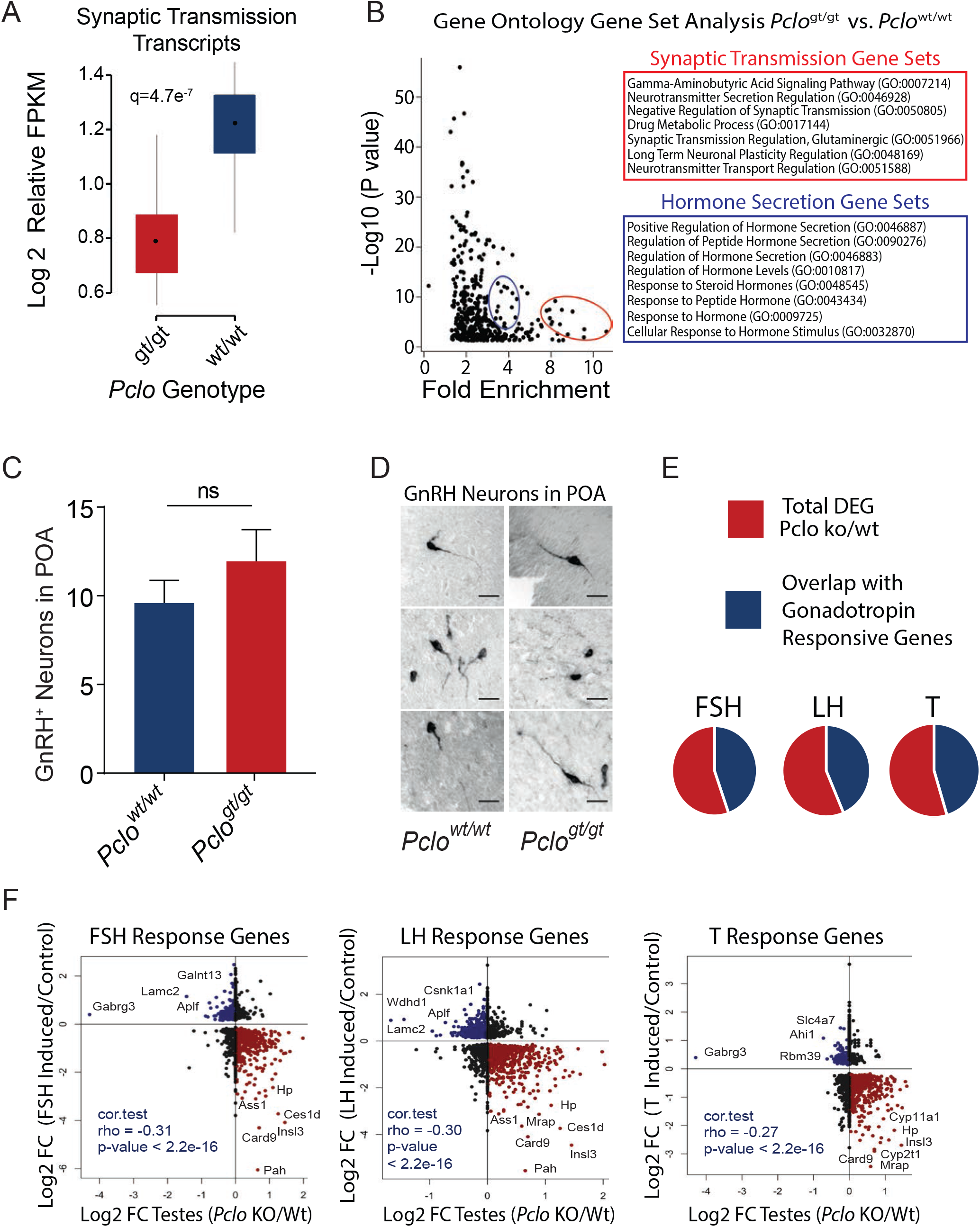
Piccolo deficiency disrupts synaptic transmission and gonadotropin responsive gene sets. (**A**) Ontology of DEGs in *Pclo*^gt/gt^ vs *Pclo*^gt/gt^ rat brains. Mean relative abundance of 80 synaptic transmission GO genes in *Pclo*^gt/gt^ vs *Pclo*^wt/wt^ rat brains (log2-fold change >1 or <-1; FDR < 0.05). (**B**) Gene Ontology gene enrichment pathway cluster analysis in *Pclo*^wt/wt^ vs *Pclo*^gt/gt^ rat brains identified misregulated clusters for Hormonal Secretion (blue circle & box) and Synaptic Transmission (Red Circle & box) GO gene sets. (**C**) Quantification of the average number of GnRH positive neurons per section within the POA, revealing that *Pclo*-deficiency does not affect the number of GnRH neurons within the preoptic area (POA). (**D**) High magnification images of GnRH positive neurons in *Pclo*^wt/wt^ vs *Pclo*^gt/gt^ rat brains (Scale bars = 10 μm). (**E**) Overlap between DEGs in *Pclo*^gt/gt^ rat testes and genes regulated by gonadotropins in rat testes[36]; FSH, Follicle Stimulating Hormone; LH, Luteinizing Hormone, T, Testosterone (log2-fold change >1 or <-1; FDR < 0.05). (**F**) Scatter plots of DEGs in *Pclo*^gt/gt^ rat testes and genes regulated by gonadotropins in rat testes [36].

### Disturbed hormonal secretion in *Pclo* mutants

In addition to downregulated *Synaptic Transmission* gene sets (**Fig 5A**), a prominent cluster of *Hormonal Secretion* gene sets was also downregulated in *Pclo*^gt/gt^ rat brains compared to *Pclo*^wt/wt^ animals (**Fig 5B**). Pathway analyses (PANTHER) further deciphered that the affected genes in *Pclo*^gt/gt^ fall most frequently into major signaling pathways of the *Gonadotropin-releasing hormone (GnRH) receptor*, followed by *Wnt, Chemokine-cytokine* and *CCKR* signaling (**S3A Fig**).

Importantly, GABA and GnRH signaling are functionally connected. During development the GABA signal, which depolarizes GnRH neurons, regulates overall GnRH neuron maturation (e.g. migration to the brain). Various GABA(A) receptor subunits are differentially expressed during the process of GnRH-1 maturation [30], and Gabra6 is a receptor subunit within embryonic GnRH-1 neurons that is replaced by Gabra2 during adult life [30]. Signaling pathways coupled to the GnRH receptor gene set control the hypothalamic-pituitary-gonadal axis that is critical for gamete development [31], and for regulating reproductive behavior [32, 33].

The reported Gabra6-positive GnRH neuronal progenitors led us to wonder whether the close-to-KO *Gabra6* phenotype in *Pclo*^gt/gt^ rats altered GnRH neuron migration patterns during development, and in turn, compromised establishment of proper GnRH receptor signaling. However, a direct assessment of GnRH neurons in the brains of adult *Pclo*^gt/gt^ rats revealed normal numbers of GnRH immuno-positive cells in the pre-optic area of the hypothalamus that projected normally into the medial eminence (**Fig 5C, D** and **S3B Fig**). Thus, GnRH neuron development within the pre-optic area of *Pclo*^gt/gt^ rats did not appear to be affected by diminished *Gabra6* expression, suggesting that other signaling mechanisms might, at least partially, compensate for *Gabra6* function during the process of GnRH neuron maturation.

Nevertheless, excitatory GABA neurons activate GnRH neurons [34] throughout adult life to affect the rate of GnRH synthesis and the pattern of GnRH release [35] as mechanisms that regulate activity of the GnRH receptor. In this context, we identify *Heterotrimeric G-proteins* as a significantly affected category in *Pclo* mutants, through which the GnRH receptor predominantly transmits its signals (**S3A Fig**). Furthermore, a major fraction of the G-protein coupled receptors (GPCRs) involved in conducting GnRH signaling on gonadotrophs are differentially enriched in the brains of *Pclo*^gt/gt^ rats compared to *Pclo*^wt/gt^ and *Pclo*^wt/wt^ rats (**S4A Fig**). Similarly, the cascade involved in mobilizing Ca^2+^ from InsP3-sensitive intracellular pools, required for the secretion of gonadotropins, is impaired in *Pclo*^gt/gt^ rats (**S4B Fig**).

Downregulation of GnRH-dependent GPCR- and Ca^2+^-stimulated processes may well affect end products of the GnRH receptor signaling pathway (e.g. luteinizing hormone, LH; follicle-stimulating hormone, FSH; **S3D Fig**). Blunted expression of GnRH signaling gene sets corresponded to reduced plasma levels of LH and FSH in the *Pclo* KO compared to WT (**S4C Fig**) Furthermore, analyzing the transcript levels of potential target genes that might be regulated by LH, FSH and/or testosterone [36] in *Pclo*^gt/gt^ vs *Pclo*^wt/wt^ rats revealed that about half of the dysregulated genes in *Pclo*^gt/gt^ testes responded to a particular hormonal stimulation in a reverse order (rho = −0.31 and p-value < 2.2e-^16^) (**Fig 5E** and **S4D Fig**). *Gabrg3, Ces1d, Card9, Insl3* and *Hp* appeared among the most affected targets of LH, FSH and/or testosterone in *Pclo*^gt/gt^ mutant rat testes (**Fig 5F**). Thus, downregulated neuroendocrine GnRH signaling failed to normally activate several gonadotropin-responsive target genes in the testis, likely contributing to the *Pclo*-deficient rat’s infertility phenotypes.

### Pclo deficiency up-regulates hypothalamic genes associated with social behavior

In addition to GABA(A) signaling, we evaluated further altered gene signatures in the *Pclo*^gt/gt^ rat brain encoding factors that would function upstream of GnRH signaling pathways (**S3A-D Fig**), potentially adding an additional layer to the complexity of the infertility phenotype. This approach identified a set of transcripts encoding neuroendocrine hormones that was selectively upregulated in *Pclo*^gt/gt^ rat brains by >3-fold vs wt (**Fig 4F** and **S5 Table**). Among the upregulated genes, *Npy, Pmch, Hcrt1, Trh* and *Avp* are known to physiologically regulate GnRH-1 neuron activity, energy balance and/or social behavior [37-40]. Collectively, the down-regulated *GnRH receptor signaling,* as well as the up-regulated *hypothalamic polypeptide hormone signaling* gene profiles in Pclo-deficient rats would suggest that Piccolo embodies a candidate presynaptic factor that regulates reproductive behavior in response to an organism’s physiological state.

### Reproductive failure in *Pclo* mutant rats is associated with neurological and behavioral defects

To follow up on altered synaptic transmission gene sets observed in the *Pclo* mutants as well as the potential behavioral aspects of the infertility phenotype, we conducted studies on brain function and behavior. Consistent with dominant GABA(A) endophenotypes (**Fig 4G**), both *Pclo*^wt/gt^ and *Pclo*^gt/gt^ mutations increased mean seizure frequencies (≥8-fold) compared to *Pclo*^wt/wt^ littermates (n=8/genotype) (**Fig 6A, left**). The EEG morphology in *Pclo*^wt/gt^ and *Pclo*^gt/gt^ rats resembled short duration absence-type seizures, displaying a characteristic 6-8 Hz spike-wave generalized onset (**Fig 6A, right**), with no convulsive activity, and functionally verifying the significance of altered *Synaptic Transmission* gene sets (**Fig 5A, B**).

**Figure 6.**
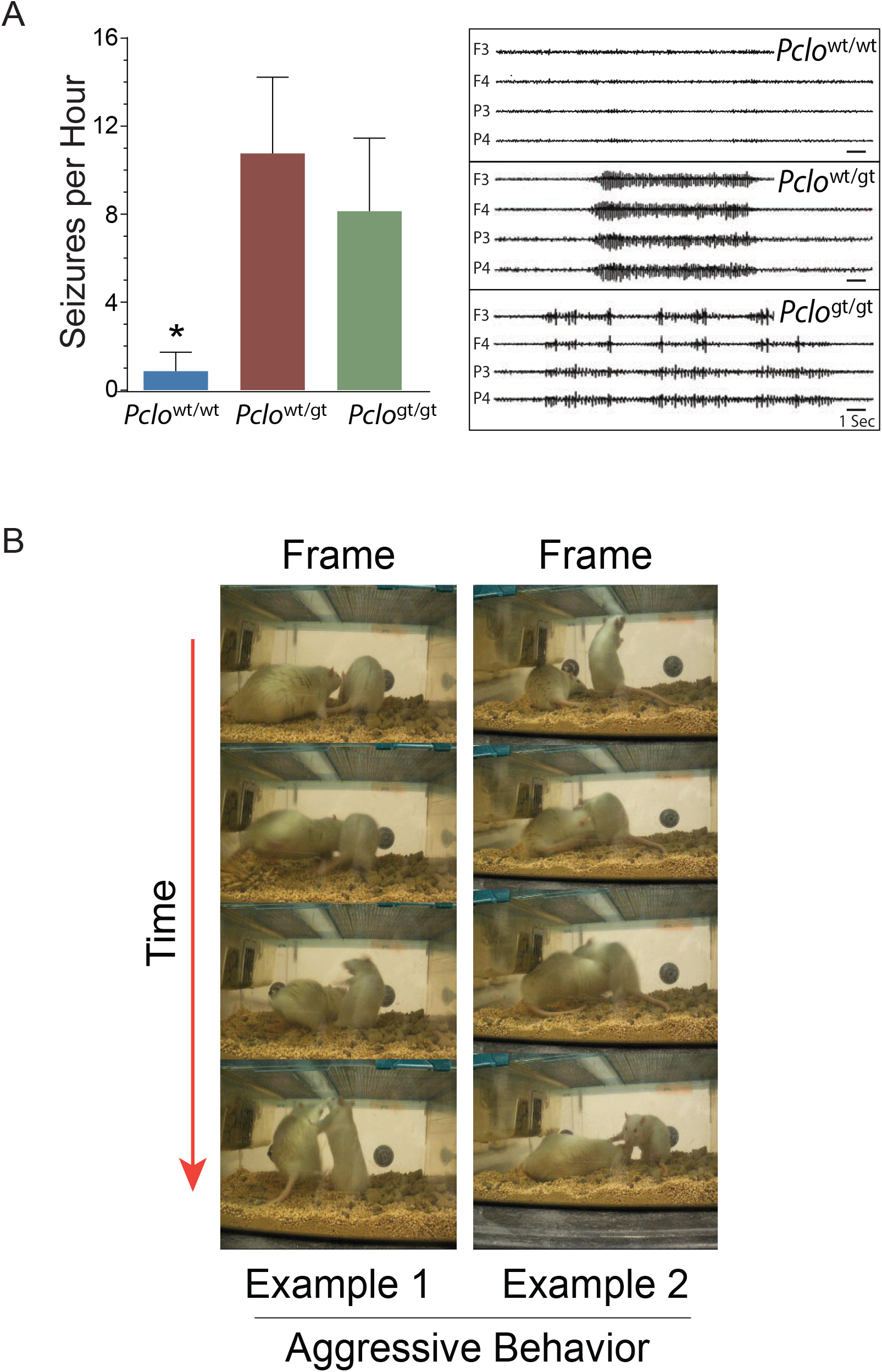
Dominant neurotransmission and recessive reproductive behavior traits in *Pclo* mutant rats. (**A**) *Top:* Seizure rates recorded in *Pclo*^wt:wt^, *Pclo* ^wt:gt^ and *Pclo*^gt:gt^ rats (n=4 rats/genotype). p<0.001 for *Pclo*^wt:wt^ vs. *Pclo*^wt:gt^ or *Pclo*^gt:gt^ rats; p=0.1 for *Pclo*^wt:gt^ vs. *Pclo*^gt:gt^ rats. *Bottom:* Representative EEG tracings recorded in *Pclo*^wt:wt^, *Pclo*^wt:gt^ and *Pclo*^gt:gt^ rat brains. (**B**) Altered social behavior in mutant *Pclo* rats. Time lapse frames showing an agitated *Pclo*^gt/gt^ male lunging defensively at a *Pclo*^wt/wt^ female rat. Similar aggressive behavior was not observed in *Pclo*^wt/gt^ or *Pclo*^wt/wt^ rats.

To find out if the Pclo mutation is associated with a behavioral phenotype, we monitored pre-copulatory social interactions that normally occur between female and male rats, including courtship, grooming and genital investigation. We observed that, in contrast to their *Pclo*^wt/wt^ littermates (**S1 Video**), female and male *Pclo*^gt/gt^ rats exhibited a relative disinterest in courting the opposite sex upon being introduced into the same cage with *Pclo*^wt/wt^ rats (p=0.0002 compared to WT littermates, n=8/genotype) (**S2 and S3 Video**). Our monitoring indicated that instinctive, normal pre-copulatory social interactions were suppressed in both female and male *Pclo*^gt/gt^ rats (**S2 and S3 Video**).

In contrast to the compatible precopulatory behavior shared between *Pclo*^wt/wt^ and/or *Pclo*^gt/wt^ rats (**S1, S4 and S5 Video**), the social phenotype displayed by *Pclo*^gt/gt^ rats of both sexes included overt aggression, biting, lunging and posturing (**Fig 6B and S2, S3, S6 Video**). Thus, in addition to the hormonal imbalance, *Pclo*-dependent neural connections function to regulate conspecific sensory responses that are likely to contribute to the infertility phenotype of *Pclo*^gt/gt^ rats. Moreover, by ~3 months of age, the aggressive phenotypes displayed by *Pclo* mutants selectively rendered the male *Pclo*^gt/gt^ rats so severely socially incompatible (**S6 Video**) that they could not be housed with male or female littermates any longer, independent of littermate genotype (n=14 of 14 male *Pclo*^gt/gt^ rats).

### Recessive *Pclo* traits are mappable to allelic markers for major depressive disorder

Mapping to a recessive phenotype, the *Synaptic Transmission* category included a severely compromised *Glutamatergic Excitation* gene set (GO:0051966, p=0.00000001) in the brain of *Pclo*^gt/gt^ rats (e.g. DEGs in the *Pclo*^gt/gt^, but not in *Pclo*^gt/wt^ mutants) (**S6 Table**). *Pclo* clustered with *Grm5, Htr2a, Negr1, Drd2, Cacna2D1* and *Dlg1* [41] as transcripts selectively down-regulated in *Pclo*^gt/gt^ rats (Cluster 1, **Fig 7A**). Among the most significantly downregulated genes were *Grm5* (Glutamate Metabotropic Receptor 5) and *Htr2a* (the serotonin [5-Hydroxytryptamine] Receptor 2A) (**S5 Table**). Both *Grm5* and *Htr2a* function as GPCRs in the signaling cascade that controls calcium mobilization and PKC activation [41, 42]. Alongside *glutamatergic* neurotransmission, *dopaminergic* neurotransmission (e.g. *Drd2*) and *Calcium signaling* (e.g. *Cacna2D1 and Dlg1*) also contributed to the recessive phenotype in *Pclo* mutants (**S6 Table**).

**Figure 7.**
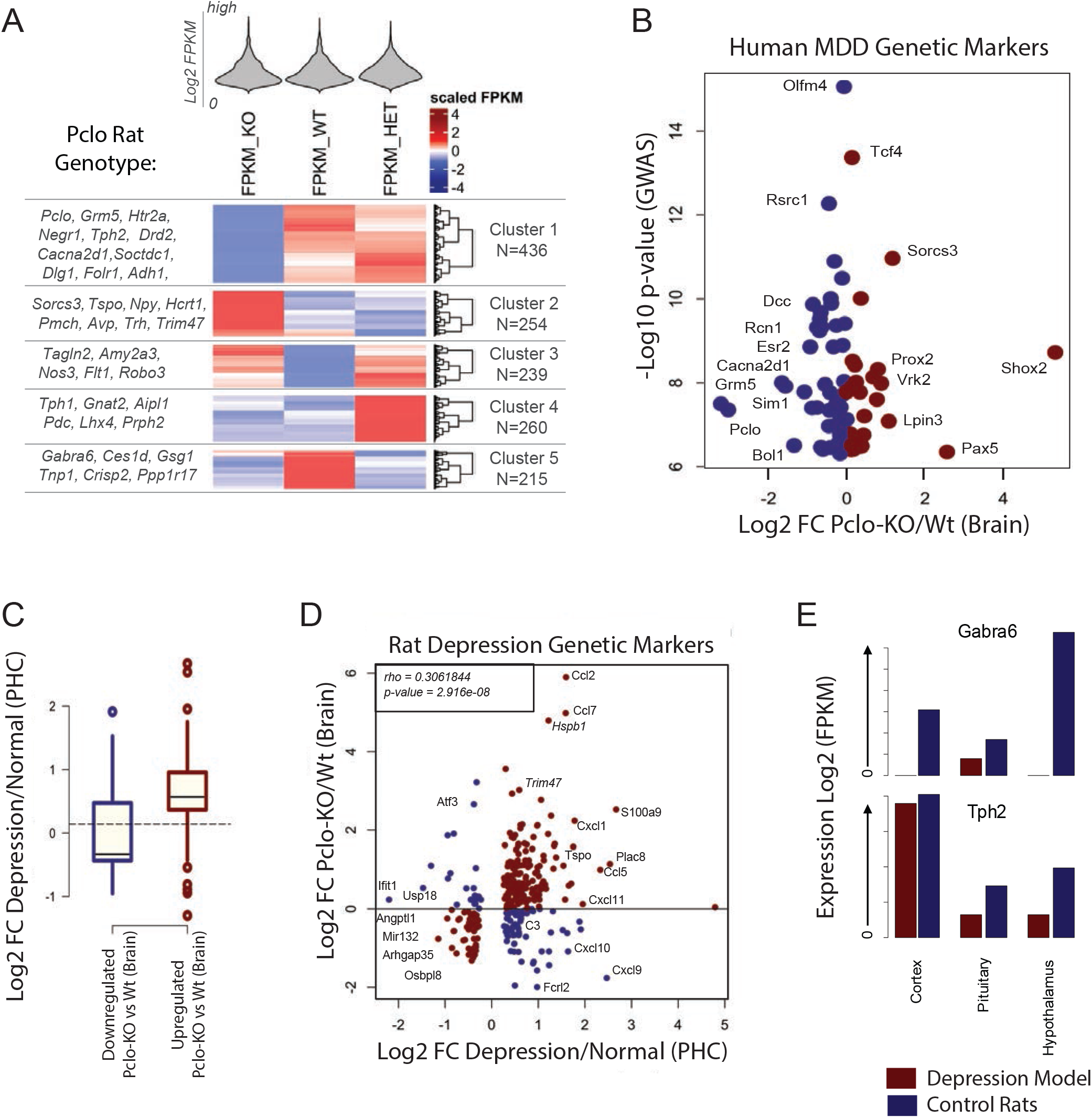
Piccolo deficiency RNA endophenotypes link to depression gene profiles. (**A**) Cluster Analysis of differentially expressed genes (DEGs) in *Pclo*^wt:wt^ (WT), *Pclo*^wt:gt^ (HET) and *Pclo*^gt:gt^ (KO) rats. (**B**) DEGs in *Pclo*^gt:gt^ vs *Pclo*^wt:wt^ (KO/Wt) rat brains are among major loci identified by human GWAS as MDD risk factors. (**C**) Dysregulated genes in *Pclo-KO* versus Wt depressed rats. Note the significance level of correlated genes. (**D**) Comparison between DEGs in depressed vs normal wt rat neurons and DEGs in *Pclo*-KO vs *Pclo*-Wt rat brains. Red dots are showing a similar pattern of expression (rho > 0.3). (**E**) *Gabra6* as prominent gene dysregulated in *Pclo-KO* rat brains is not detected in Pituitary and Hypothalamus of depressed rats (*top*). Depression-related marker *Tph2* shown in comparison (*bottom*).

Intriguingly, 6 dysregulated genes in the recessive *Pclo* traits (e.g. *Pclo*, *Grm5*, *Cacna2D1, Negr1, Sorcs3* and *Drd2*) are among the recently reported 44 key risk factors of major depressive disorder (MDD), identified by a human genome-wide association study containing 135,458 MDD cases and 334,901 controls [43] (**Fig 7B**). To test for a potential association between the biological processes dysregulated in *Pclo*^gt/gt^ rats and a neurological disorder, we data-mined and compared the transcriptome of brain samples derived from an MDD rat model [44] and our *Pclo*^gt/gt^ rat (**Fig 7C**). Our strategy identified a robust list of 408 genes that were similarly affected in both models (*rho = 0.306* and *p-value = 2.916e-08)* (**Fig 7D**), supporting a transcriptome-level relationship between the biological processes dysregulated in *Pclo*^gt/gt^ rats and a rat model of MDD. Notably, the shared list of MDD transcripts encoded genes that have been associated with various features of depression, such as cortical dementia (e.g. *Trim47*), moodiness (e.g. *S100A9*), enhanced microglial activation (e.g. *Tspo*) and depression followed by immune challenge (e.g. **Fig 7D**). Notably, *Gabra6*, among the most highly dysregulated genes in *Pclo* KO rats, was also not detectable in the hypothalamus and cortex samples of the MDD rat model (**Fig 7E**), connecting *Gabra6* with both *Pclo* and MDD phenotypes [20, 45].

### Cross-species analysis reveals robust differences in rat reproduction mutant phenotypes

Finally, we compared our rat phenotypes to data recorded in other species harboring loss-of-function mutations in orthologous genes (**S4 Table**). Nine mutated rat genes (*Atg13, Btrc, Dlg1, Grik3, Pclo, Slc1a3, Spaca6, Zmynd8*, and *Ube2k*) have mutated orthologs in mice [Mouse Genome Informatics (MGI), the International Mouse Phenotype Consortium (IMPC) and the National Center for Biological Information (NCBI) databases], whereas 5 of our mutated orthologs have been characterized in plants, yeast, worms, flies or frogs (*Alk3, Atg13, Btrc, Dlg1, Pan3*) (**S4 Table**). Strikingly, the ‘shortened life span’ caused by mutations in *Atg13* has been reported across multiple species including plants, yeast, worms, flies, mice and rats (**S4 Table**).

In humans, genome-wide association studies (GWAS) have implicated orthologs for 12 of the mutant rat genes we analyzed (*Abca13, Alk3, Atg13, Btrc, Dlg1, Exoc6b, Fstl5, Gsgl1, Grik3, Pclo, Slc1a3, Ube2q2*) as either risk factors or candidate risk factors for various human disease processes (**S4 Table**). About half of these reported disease factors are associated with neurological/behavioral disorders (e. g. *Abca13, Dlg1, Exoc6b, Grik3, Pclo, Slc1a3*) (**S4 Table**).

Of note, *Dlg1*^gt/wt^ rats displayed reduced fecundity and altered behavior (**S4 Table**). Intriguingly, a *Dlg1*-null mutation was reported to disrupt courtship and mating in flies [46]. *Dlg1*^gt/wt^ rats were consistently observed in back cage corners, remained socially isolated following pairing with respective wildtype males, and only produced offspring on one occasion (n=4 pups; 3 wildtype males and 1 mutant female) after pairing for extended periods (>10 months) with different wildtype males and appeared otherwise healthy (See **S4 Table**). In *Pclo*^gt/gt^ rat brains, *Dlg1* (a.k.a. *Synapse-Associated Protein 97* or *SAP97*) was observed as the most downregulated gene in in the GO category of Calcium Signaling (**S4B Fig**), unveiling a potential connection between *Pclo* and *Dlg1* to regulate conspecific social behavior.

Our comparative analysis provided several additional examples where gene mutations analyzed here in rats produced significantly different phenotypes in another species with orthologous gene mutations (S4 Table). Such differences may reflect the quality of the knockout and/or species-dependent differences in biology. As a prime example, while fertility and behavior is normal in *Pclo-*deficient mice [47], our *Pclo* mutants were infertile, complicated with social incompatibility, and a potential genetic link with depression (Figs 4A, 7B-E and S2, S3, S6 Videos).

## Discussion

Here, we characterize a pool of 12 distinct mutant rat strains that are unable to reproduce (*Alk3, Atg13, Dlg1, Btrc, Exoc6b, Pan3, Pclo, Slc1a3, Spaca6, Tmx4, Ube2k, Zmynd8*). The mutant rat strain pool was derived from a library of recombinant spermatogonial stem cells harboring randomly inserted *Sleeping Beauty* gene trap transposons [17]. The reproduction phenotypes we identified in rats were all associated with different steps in spermatogenesis or embryonic lethality except for three mutant strains (*Atg13, Dlg1, Pclo*). Of the later mutants, *Atg13* and *Pclo* strains stood out by exhibiting a “complex” phenotype that allowed us to decipher novel aspects of reproduction.

Our *Atg13*^gt/gt^ rats displayed abnormal autophagy markers, gross renal abnormalities and inflammation-like phenotypes that preceded death in early adulthood. Atg13 (Autophagy related 13) is the master metabolic sensor for toggling between AMPK1-dependent cellular torpor (i.e. autophagy) and ULK1-repressed mTORC1-dependent cell growth. The *Atg13*^gt/gt^ rat phenotype might be related to the loss of a phylogenetically conserved Ulk1-binding peptide encoded by *Atg13’s* terminal exon (**Fig 3A**). *Atg13’s* COOH-terminus has been implicated in activating the main autophagy-initiating complex [48]. Similar to the rat *Atg13*^gt/gt^ phenotype, dysfunctions in *Atg13* have been associated with nephrological/immunological problems and autophagy in humans [49, 50]. Mice harboring either a frameshift mutation in *Atg13* exon 5 or a gene trap in *Atg13* exon 1, by contrast, exhibit a more severe phenotype and die *in utero* due to heart defects [51]. Notably, the end-stage pathology of *Atg13*^gt/gt^ rats correlated with immotile, degenerating caudal epididymal spermatozoa, likely associated with the premature aging phenotype. While the *Atg13*^gt/gt^ rat represents an excellent model to study the connection between premature aging, fitness and fertility, our *Pclo* and *Dlg1* mutants highlighted how traits linked to human neurological disorders might disrupt rat reproductive behavior.

Curiously, the *Pclo*^SBΔ4-25^ mutation disrupted reproduction, but induced more “global” changes in the brain transcriptome than in the testis, suggesting a possible crosstalk between the brain and gonads. The most significant changes in both tissues affected GABAergic signaling via GABA(A) receptors. Our data support a scenario, where the infertility phenotype is connected to the altered composition of GABA(A) receptor subunits associated with the GnRH signaling cascade. GABA has been shown to play an important role in the maturation of gonadotrophin-releasing hormone (GnRH)-1 neurons during development and in regulating the pulsatile release of GnRH in adults [30, 34, 35]. Accordingly, impaired GnRH receptor signaling would translate into reduced responsiveness of testicular target genes (**Fig 5E-G**). The most significantly dysregulated target gene in Pclo^*gt/g*t^ rat testes also encodes a GABA(A) receptor, *Gabrg3, gamma subunit 3* (**Fig 4G**), suggesting that a crosstalk between brain and testes may also involve a mechanism that regulates *Gabrg3*-dependent GABAergic tone. Thus, our *Pclo*^SBΔ4-25^ mutant rat model holds potential to help address the long-standing debates on how GABAergic tone in the brain and testes is functionally linked to *GnRH neuron receptor activity* [52] and reproductive behavior [32, 33].

The *Pclo*^SBΔ4-25^ rat model exhibited additional GABAergic neuropathies. Both homozygous and heterozygous *Pclo*^SBΔ4-25^ rats develop generalized seizures (**Fig 6A**), similar to seizures observed in children homozygous for *Pclo*^Δ6-stop^ of *pontocerebellar hypoplasia type 3a* [18]. Disturbed *GABAergic synaptic transmission* in *Pclo* mutants likely affects the balance between inhibition and excitation and thereby provokes seizures, manifesting itself as epileptiform activity [34, 35]. *Pontocerebellar hypoplasia type 3a* has been also connected with impaired recycling of synaptic vesicles regulated by Pclo [53, 54].

Intriguingly, among the differentially expressed genes in *Pclo*^SBΔ4-25^ rats, we found an association with the transcriptomes in brains of rats modeling depressive disorders (**Fig 7B-E**). The top candidates of depression-related genes identified here as DEGs in *Pclo* rats^SBΔ4-25^ were involved in *glutamatergic* and *dopaminergic* neurotransmission and *neuronal calcium signaling* pathways, and further matched key allelic neurological markers identified independently in large scale GWAS cohorts of humans diagnosed with MDD (e.g. *Pclo, Grm5, Htr2a, Sorcs3, Negr1, Drd2*) [43, 55]. Affective disorder and limbic system neurotransmission phenotypes reported in MDD patients harboring *Pclo* variants were previously demonstrated to disrupt emotional processing in response to conspecific facial cues [56, 57]. By analogy, the pre-copulatory mating behavior and aggression phenotypes reported here in *Pclo*^SBΔ4-25^ rats demonstrate Piccolo’s potential control over sensory responses to social cues.

Importantly, while both *Piccolo* and *Gabra6* variants are associated with MDD and altered emotional processing based on studies in humans [20, 45], no direct sexual connection to either gene has been reported in humans. The functional significance of the tight control of *Gabra6* expression by *Pclo* has yet to be investigated, but one possibility is that a loss of synaptic integrity leads to *Gabra6* down-regulation [58]. Even so, reports on *Gabra6* KO mice suggest that they exhibit no behavioral phenotypes [59, 60], indicating that the complexity of phenotypes observed in *Pclo*^gt/gt^ rats may not be entirely explained by *Gabra6*-deficiency alone.

In contrast to neurological phenotypes caused by *Pclo* variants in rats reported here, mice that lack the full calcium sensing coil-coil domain encoded by *Pclo* exon 14 (*Pclo*^Δ14^ mice) behave normal and are fertile [47, 61]. While we did not observe a significant difference in homozygous *Pclo*^SBΔ4-25^ rat body weights (**S3 Table**), homozygous *Pclo*^Δ14^ mice displayed reduced body weights and enhanced postnatal mortality, consistent with a negative energy balance [47]. In contrast, *Pclo*^SBΔ4-25^ rats, curiously, like *Dlg1-*deficient flies [46], exhibit reproduction abnormalities that may be attributed to suppressed pre-copulatory and copulatory behavior. When compared to *Pclo*-dependent sensory responses in humans [56, 57], *Pclo*-dependent reproductive behavior displayed by *Pclo*^SBΔ4-25^ rats might point to compromised synaptic transmission in the olfactory system, limbic system and/or hypothalamus as brain regions impacted by *Pclo* deficiency.

In summary, by forward genetics in rats, we annotated *Pclo* as a candidate reproductive factor that controls behavioral responses to sensory input. Studies can now be aimed at defining how Pclo-dependent neural circuits in the rat control social behavior, and prospectively, how Pclo-dependent neuroendocrine signaling integrates social responses with metabolic state.

## Materials and Methods

### Mutant rat strains

Mutant rat strains harboring *Sleeping Beauty Beta-Geo* gene trap transposons were originally transmitted to F1 progeny from a donor recombinant spermatogonial stem cell library [17]. Recipient males were bred with wildtype females to produce a panel of mutant rat strains that harbor gene traps within protein coding genes [17]. Eighteen heterozygous F1 mutant rat strains (**S1 Fig, S1 Table**) derived from an original pool of >150 *Sleeping Beauty β-Geo* gene trap strains (Cryopreserved at UT Southwestern’s Mutant Rat Resources: SSCLBR Cat & RRID) were maintained live due to an expressed interest in and/or requests for respective strains by researchers representing a broad spectrum of biomedical fields (**S2 Table**). Rat protocols were approved by an Institutional Animal Care and Use Committee (IACUC) at the University of Texas Southwestern Medical Center, as certified by the Association for Assessment and Accreditation of Laboratory Animal Care International (AAALAC).

### Rat breeding for forward screen

A panel of 18 heterozygous F1 *Sleeping Beauty* gene trap mutant rat strains was evaluated for their ability to reproduce (**S2 Table**). Based on a ~11 kb deletion from mouse Chr17 containing *Spaca6* and *Has1* that blocked sperm-egg fusion in mice [21], the *Spaca6*^gt/gt^ mutant rats were included in the current study to provide a control gene trap hypothesized to disrupt reproduction. Additionally, *Rgs22*^gt/gt^ mutant rats were included in the current study to provide a control gene trap hypothesized not to disrupt reproduction, due the gene trap cassette being inserted in the 3’ to 5’ orientation.

Founder-derived F1 mutant progeny were crossed with wildtype rats to produce F2 mutants. Males and females for 17 of 18 F2 heterozygous mutant strains successfully produced littles, of which, mean litter sizes produced by 15 of the F2 heterozygous mutant strains were comparable in size to wildtype Harlan, Sprague Dawley rat stocks (**S2 Table**). Only *Dlg1*^wt/gt^ females were identified as sub-fertile after pairing heterozygotes with wildtype rats of opposite sex for >10 months. One *Dlg1*^wt/gt^ female produced a single mutant female in one total litter (n=4 pups); however, the second generation *Dlg1*^wt/gt^ female failed to reproduce litters after subsequent pairings with fertile males for 12 months.

Male and female (F3) heterozygous mutants from the other 17 strains were generated from separately outbred parents (Harlan, SD) and paired at 3-4 months of age to generate F4 homozygous mutants. Heterozygous mutant pairs that produced litters and displayed markedly reduced Mendelian rates towards generation of homozygous mutant progeny were classified as embryonic lethal (i.e. no homozygous mutant F4 progeny; n>50 total pups/strain except for *Alk3*^wt/gt^ mutants, where n=35). Viable F4 homozygous mutants were paired with proven wildtype breeders (Harlan, SD) of opposite sex between 3-4 months of age to identify recessive mutations that transmitted significant changes in mean litter size. If F4 homozygotes failed to generate progeny by 3-4 months after pairing with a wildtype breeder, they were paired with a second wildtype proven breeder from Harlan, SD. Genes were classified as required for rat reproductive success under our standard housing conditions if homozygous mutations blocked multiple F4 progeny (n=2-4 homozygous mutant breeders/sex) from producing any offspring after pairing with 2 consecutive wildtype proven breeders of similar age over a span of >10 months. Adult lethal homozygous *Atg13* mutants demonstrated health decline between 3-4 months of age (i.e. shortly after setting up breeder pairs).

### Genotyping mutant rat progeny

Endogenous gene-specific PCR primers near *Sleeping Beauty* integration sites were used in combination with transposon-specific primers to genotype progeny from familial generations F1 and F2 for newly generated mutant rat lines. Genomic sites of transposon integration were defined in F1 progeny by splinkerette PCR[17] and sequence analysis alignment on genome build RGSC v3.4 (Rn4). Genotyping results were verified by Southern blot hybridization assays of genomic DNA digested with XmnI and XbaI using a probe specific for the *EGFP* transgene and the *LacZ* portion of the *β-Geo* insert in the *Sleeping Beauty* transposon[17]. Restriction analysis by Southern blot estimated ~7 transposon integrations/stem cell genome, which following random segregation and ploidy reduction during meiosis yielded ~3.5 transposon integrations/donor-derived spermatozoa,or founder-derived mutant F1 pup[17]. Phenotypes in *Atg13, Btrc, Pclo, Pan3, Spaca6* and *Ube2k Sleeping Beauty* mutant rat strains were analyzed in F4 animals produced from F3 breeder pairs harboring only their respective, *Sleeping Beauty* transposon integration (i.e. single copy gene trap transposon F3 mutants).

### Phenotype database and literature analysis

European Conditional Mouse Mutagenesis Programme (EUCOMM), Knockout Mouse Project (KOMP), Mouse Genome Informatics (MGI), International Mouse Phenotype Consortium (IMPC) and National Center for Biological Information (NCBI) databases provided records on mouse gene orthologs. NCBI PubMed, Gene and the Rat Genome Database (RGD) provided records on rat gene orthologs. Human phenotypes for mutant orthologs were searched in publicly available NCBI Genetics and Medicine databases, including: PubMed, Gene, Online Mendelian Inheritance in Man (OMIM), Database of Genotypes and Phenotypes (dbGaP); and the National Human Genome Research Institute’s Catalog of Published Genome Wide Association Studies (NHGRI GWAS Catalog). NCBI PubMed and Gene were searched to identify phenotypes available for *Arabidopsis, Saccharomyces, Caenorhabditis, Drosophila, Danio* and *Xenopus* species. PhenomicDB database verified results from above database searches across all species. Literature comparisons for phenotypes caused by mutations in rat and mouse orthologs published independent of the current study are summarized in **S4 Table**. Embryonic lethality or postnatal lethality prior to reproductive age was categorized as blocking reproduction. Fishers Exact t-test (two-tailed) was used to analyze phenotypic proportions of viable versus sub-viable, viable versus embryonic lethal, fertile versus infertile, mating versus non-mating.

### Sperm counts and copulation

Epididymides were harvested from adult rats between 120-180 days of age and dissected free of surrounding fat and connective tissue for measuring weights, counting spermatozoa and histological analysis. To estimate spermatozoa numbers/rat, each epididymal caput and cauda were dissected apart from the corpus and separately placed into 3.8 cm^2^ wells of a 12 well plate containing 1.5 ml DHF12 nutrient media [Dulbecco’s Modified Eagles Medium:Ham’s F12 (1:1); Sigma, D8437] 1x antibiotic antimycotic solution (Invitrogen, cat. no. 15240-062). Spermatozoa were released by thoroughly mincing each epididymal piece for 30 sec and allowing the spermatozoa to disperse into the medium for 25 min. Large pieces of epididymal tissue were removed with forceps and discarded. One ml of the epididymal cell-containing medium was carefully filtered through a 100 μm cell strainer (BD Biosciences, Inc.) into a 1.5 ml microfuge tube prior to counting using a Hemocytometer chamber. To assess breeding behavior and detect copulation, rats were paired with a single wildtype mate just prior to the end of the daily light cycle (4:00-5:00 pm central standard time). The following morning (7:00-8:00 am central standard time), each female was examined for the presence of spermatozoa in the vagina. A foam swab tip was used to collect a vaginal smear, which was then analyzed by phase contrast microscopy to detect presence of sperm.

### Analysis of RNA-seq data from Pclo rats

Single end 100 bp RNA-seq libraries were prepared from brain, liver and testis tissues of ~6-month-old Pclo^gt/gt^, Pclo^gt/wt^, Pclo^wt/wt^ rats. The libraries were run on *Illumina Hiseq 2000* sequencer (Total number of reads was ~550-600 million). For *basecalling* we used the *Illumina Casava1.7* software. Reads were than aligned to the reference human genome version *rn6* by using *Tophat2/bowtie2*. This approach has provided a *refseq_rn6* gene model that guided the assembly process of the transcriptome. We checked the quality of the sequencing and the mapping by *Fastqc* and by *RNASeqQC*, repectively. Due to the negligible technical variances, the read counts of a gene had a Poisson distribution, thus we could apply the single-replicate model to analyze the data. We calculated Read counts using *featureCounts* from the *subread package* (http://subread.sourceforge.net/). Fragments Per Kiolobase of RNA per Million mapped reads (FPKM) was calculated using *bamutils* (http://ngsutils.org/modules/bamutils/count/).

### Analysis of differentially expressed genes

Random Variable1 (*Var1*) = *n.l.x*, where *x* (Random Variable2) is the expression level of a gene (e.g., in RPKM (Reads Per Kilo bases per Million reads) *n* is reflecting the sequencing depth and *l* is the gene length. The method proposed by Anders and Huber was used to calculate *n*[62]. To generate more robust and accurate *Fold change* values from unreplicated RNA-seq data, we determined the normalization constant and variance by pasting the two random variables in the published algorithm of: (http://bioinformatics.oxfordjournals.org/content/early/2012/08/23/bioinformatics.bts515.full.pdf+html). To identify the Gene Ontology (GO) categories that were overrepresented in the Piccolo mutants, we compared samples from the brain and testis of *Pclo*^gt/gt^ and *Pclo*^wt/gt^ vs *Pclo*^wt/wt^ rats, with the entire set of rat genes as a background.

### Electroencephalogram (EEG) recording and analysis

Twelve adult rats (6 males, 6 females) were surgically prepared for EEG experiments with 4 rats in each experimental group (*Pclo*^wt/wt^, *Pclo*^wt/gt^, *Pclo*^gt/gt^). Rats were anesthetized using a gas anesthesia machine with ~3% isoflurane in a 1 L/min mixture of 70% nitrous oxide and 30% oxygen. Four epidural recording electrodes made from #00-90 x 1/8 inch stainless steel screws were placed at the following stereotaxic coordinates: A-P ±2.0 mm, lateral ±3.0 mm and A-P - 4.0 mm, lateral ±3.0 mm along with a reference and ground screw over the olfactory bulb and cerebellum, respectively. Electrodes were attached by a flexible wire (kynar, 30 ga) to a custom 6-pin microconnector (Omnetics) and secured with dental acrylic. Rats received the analgesic buprenorphine (0.05 mg/kg) as necessary following surgery and were allowed to recover for at least 7 days prior to any experimentation. Following recovery from electrode implantation, each rat was placed in a custom acrylic recording cage (Marsh Designs, Peoria, AZ) and connected to a Tucker-Davis Technologies (Alachua, FL) RZ2/PZ3 neurophysiology workstation through a flexible cable suspended from the top of the cage with an interposed commutator to allow rats free access to food and water without twisting the cable. Continuous video/EEG (300 Hz sampling) was recorded for each rat simultaneously for 7 days and read by a user blinded to the experimental grouping for the presence of seizures and epileptiform activity. Seizure activity was marked at the beginning and end of each event to account for seizure duration, and the numbers of seizures per hour were calculated.

### Western blot analysis

To analyze Piccolo expression, brains were dissected from wildtype, heterozygous mutant, and homozygous mutant Sprague Dawley rats and homogenized in 1.5 ml/0.5g tissue, ice-cold lysis buffer (50 mM HEPES, pH 8.0, 150 mM NaCl, 1 mM EDTA, 10% glycerol, 1% Triton X-100, 10 μg/ml aprotinin, 10 μg/ml leupeptin and 1 protease inhibitor Data Sett/12.5 ml) for 30s using a PTA-7 probe, setting 5, PT10-35 polytron (Kinematica). The homogenates were incubated on ice for 15–20 min and then centrifuged at 3000x*g* for 10 min at 4°C in a GPR Data Settop centrifuge (Beckman, Inc.). The supernatant solutions were centrifuged at 15,800x*g* for 15 min at 4°C in a microcentrifuge (Model 5042, Eppendorf, Inc.) and the resultant supernatant fractions were stored at −80°C. 160 μg of protein was separated on 4-15% Mini-Protean TGX gels (BioRad, Inc.), and then transferred to nitrocellulose. Samples were not heated prior to loading. Nonspecific, protein binding sites were blocked by incubating membranes overnight at 4°C in blocking buffer: TBST (Tris-buffered saline with Tween-20: 10 mM Tris–HCl, pH 7.5, 150 mM NaCl, 0.1% Tween-20) containing 5% nonfat dry milk. Membranes were washed three times in TBST and incubated for 1 h at 22–24°C using rabbit anti-Piccolo (Synaptic Systems cat. no. 142002) diluted 1:2000 in blocking buffer. Membranes were washed three times in TBST (0.3% Tween-20) and incubated 45 min, 22-24ºC with peroxidase-conjugated, anti-rabbit IgG (Jackson Immunoresearch) diluted 1:50,000 in blocking buffer. Membranes were washed three times in TBST and protein bands detected using the enhanced chemiluminescence detection method (ECL, Amersham, Inc.). Blots were stripped and re-probed with 1:20,000 dilution of mouse anti-TUBA1a (MU-121-UC, Biogenex, Inc.).

Rat embryonic fibroblast (REF) cultures were extracted in RIPA buffer (50 mM Tris pH 7.4, 150 mM sodium chloride, 1 mM EDTA, 1% IPEGAL, 0.25% deoxycholic acid) plus protease inhibitor and phosphatase inhibitor Data Setts (Roche Applied Science). 11 μg protein was separated on NuPAGE 4-12% Bis-Tris gels (Invitrogen, Inc.) and then transferred to nitrocellulose membranes. Nonspecific protein binding sites were blocked by incubating membranes overnight at 4°C in blocking buffer: TBS (Tris-buffered saline: 10 mM Tris-HCl, pH 7.4, 150 mM NaCl) containing 1X Western Blocking Reagent (Roche Applied Science, Inc.). Antibodies were diluted in TBS containing 0.5X Western Blocking Reagent + 0.1%Tween-20. Membranes were incubated in primary antibody for 1-2 hours at 22-24°C. Membranes were washed 4 x 5 min in TBST (0.1%-0.3% Tween-20), incubated in IRDye secondary antibody for 45-60 min, washed again 4 x 5 min, and scanned on an Odyssey Classic Quantitative Fluorescence Imaging System, Model 9120, Licor Biosciences, Inc. Images were analyzed with Odyssey software version 3.0.21. Primary antibodies: Rabbit anti-LC3A from Cell Signaling Technology, Inc, #4599, 1:300; Mouse anti-Atg13 from Medical and Biological Laboratories, Ltd, #M183-3, 1:1000; Guinea pig Anti-p62 from Medical and Biological Laboratories, Ltd, #PM066., 1:2000. Secondary antibodies were all from Licor Biosciences: Goat anti-rabbit IRDye 800CW #926-32211, 1:15000; Goat anti-mouse IRdye 680LT 1:20000; Donkey anti-guinea pig IRDye 800CW #926-32411, 1:15000.

### ELISA on Rat Plasma

Plasma LH and FSH levels were measured using ELISA Kits from CUSABIO according to the manufacturer’s instructions (Rat FSH Cat# CSB-E06869R; Rat LH Cat# CSB-E12654r from CUSABIO).

### Histological sectioning and staining

Hematoxylin/Eosin (H&E), periodic acid-Schiff’s (PAS) and Trichrome staining on histological sections from rat tissues were conducted by standard procedures at the Molecular Pathology Core Laboratory, UT Southwestern Medical Center in Dallas.

### Preparing frozen sections

To prepare frozen testis sections for labeling with antibodies, testes were dissected from rats, perforated by puncturing three equally spaced holes in the *tunica albuginea* along each longitudinal axis of the testis using a 27 gauge needle, and fixed for ~18 hr at 4°C in 0.1M sodium phosphate buffer, pH 7.2, containing 4% paraformaldehyde. Fixed testes were equilibrated through a 10%, 18% and 25% sucrose [wt/v, dissolved in 1x phosphate buffered saline (PBS; Invitrogen Inc, cat no. 14040-182)] gradient by sequential overnight incubations (~24 hr) at 4°C in 20 ml of each respective sucrose solution. Once equilibrated to 25% sucrose, testes were embedded in tissue freezing medium (Electron Microscopy Sciences Inc., #72592) and frozen using a Shandon Lipshaw (#45972) cryo-bath. Frozen testes were used to prepare a parallel series of 8 μm cryo-sections. Frozen sections were stored at −40°C until use in immunofluorescence assays as described below.

### Fluorescence immunohistochemistry

Prior to labeling studies, sections were equilibrated in air to ~22-24ºC for 15 min, hydrated in Dulbecco’s phosphate-buffered saline (PBS) (Sigma, D8537) at 22-24ºC for 10 min, heat-treated at 80°C for 8 minutes in 10 mM sodium citrate (pH 6.0) and then incubated for 1 hr at 22-24ºC in blocking buffer [Roche Blocking Reagent (1% v/v) diluted in 0.1M Sodium phosphate buffer, containing Triton X100 (0.1% v/v)]. Sections were then treated for 18-24 hr at 22-24ºC with respective antibodies diluted in blocking buffer at the following concentrations: [1:400 mouse anti-Sall4 IgG (H00057167-M03, Abnova, Inc); 1:400 rabbit anti-phospho-H2A.X (Ser139) IgG (07-164, Millipore, Inc); 1:400 rabbit anti-phospho-Histone H3 (ser10) IgG (06-570, Millipore, Inc)] diluted into Roche blocking (1% w/v) reagent. After treatment with primary antibodies, sections were washed 3 times for 10 min/wash in 50 ml PBS and then incubated for 40 min at 22-24ºC with respective AlexaFluor594 (Invitrogen, Inc), or AlexaFluor488 (Invitrogen, Inc) secondary antibodies diluted to 4 μg/ml in PBS containing 5 μg/ml Hoechst 33342 dye (Molecular probes, cat no. H3570). After treatment with secondary antibodies, sections were washed 3 times at 10 min/wash in 50 ml PBS. After the 3^rd^ wash in PBS, sections were cover-slipped for viewing using Fluorogel mounting medium (Electron Microscopy sciences, cat no. 17985-10). Images were acquired using an IX70 Olympus fluorescence microscope (Olympus Inc.) equipped with Simple-PCI imaging software (C-Imaging Systems, Compix, Cranberry Township, PA).

### Rat embryonic fibroblast culture

Primary rat embryonic fibroblast (REF) cultures were prepared from E14.5 embryos dissected from wildtype female rats after mating with *Atg13*^wt/gt^ male rats. Timed mating was established as described above in the section on *Sperm Counts and Copulation*. Uteri were dissected from pregnant females and washed with 10 ml DHF12 medium, 1% Penicillin-Streptomycin solution (v/v). The heads and visceral tissue were removed from each isolated embryo. Visceral tissue was discarded. Tissue from the upper portion of the head was used to isolate genomic DNA and genotype embryos for the *Atg13* gene trap mutation. The remaining thoracic portion was washed in fresh DHF12 medium, transferred into tubes containing 5 ml 0.05% trypsin/1mM EDTA solution, minced for 2 minutes and then incubated at 37°C for 20 min. After incubation, REF culture medium [DMEM (Sigma, D5648-10XL), 10% fetal bovine serum (Tissue Culture Biologicals, 104300), 1% Penicillin/Streptomycin (Hyclone, SV30010)] was added to the cell suspension and the cells were dissociated further by gentle trituration (5 strokes) using a p1000 Eppendorf tip. The cell suspension was centrifuged 4 min at 120 x *g* and the supernatant was discarded. The cellular pellet was retained, suspended to 15 ml in fresh REF medium, plated into 10cm plastic tissue culture dishes (Corning, Inc.) and then incubated at 37°C, 5% CO2 overnight. REFs were fed 15 ml fresh medium every 48 hrs, and sub-cultured using the 0.05% trypsin/1mM EDTA solution to harvest attached cells from culture dishes every 2-3 days. Harvested REFs were passaged by plating at ~10^4^ cells/cm^2^ in 3 ml/cm^2^ REF medium. REF cultures were maintained at 37°C, 5% CO2, and used for experiments at passage 4. REFs were treated for 24 hr with or without 3 mM ammonium chloride (Fluka, 09718), 100 nM Rapamycin A (LC Laboratories, R-5000) and, or 3 nM Bafilomycin A1 (Sigma, B1793) prior to preparing lysates for western blots.

### Perfusion, Sectioning and Immunohistochemistry of rat brains

*Perfusion:* Adult rats (P100) were first sedated in Isoflurane (Abbott GmbH & Co. KG, Wiesbaden, Germany) and then deeply anesthetized with a mix of 20 mg/ml Xylavet (CO-pharma, Burgdorf, Germany), 100 mg/ml Ketamin (Inresa Arzneimittel GmbH, Freiburg, Germany) in 0.9% NaCl (B/BRAUN, Melsungen, Germany). Afterwards the heart was made accessible by opening the thoracic cavity, and a needle inserted into the left ventricle and the atrium cut open with a small scissor. Animals were initially perfused with PBS and then with freshly made 4 % PFA, before dissecting and further incubated for 24h in 4 % PFA at 4°C. Brains were then cryoprotected in 15% and then 30% sucrose at 4°C for 24h. Brains were then frozen using 2-methylbutane (#3927.1, Carl-Roth, Karlsruhe, Germany) cooled with dry ice to −50°C and stored at −20°C. *Brain sectioning*. 20 μm thin serial sections were cut from frozen brains using a cryostat (Leica Mikrosysteme Vertrieb GmbH, Wetzlar, Germany). Slices transferred to a microscope slide (Superfrost Plus, #H867.1, Gerhard Menzel B.V. & Co. KG, Braunschweig, Germany), dried at RT for at least 1h and stored at - 20°C. *Immunohistochemistry*. 3,3’-Diaminobenzidine (DAB) staining of 20 μm coronal brain sections labeled with mouse anti GnRH antibody performed as previous described (Brinschwitz et al., 2010). In brief, thawed sections were dried for 30 min at RT and washed 3x for 10 min in PBS-T (PBS 1X (Thermo Fisher Scientific, Waltham, USA) + 0.025% Triton X-100 (#3051.2, Carl-Roth, Karlsruhe, Germany) and endogenous peroxidase was blocked for 10 min with 0.3% H2O2 in PBS, before blocking for 2h at RT in blocking solution (PBS plus 10% normal goat serum and 1% BSA). Sections were then incubated in primary mouse anti GnRH antibody (1:500, HU4H, provided by H. Urbanski, Oregon Regional Primary Center, Beaverton, OR) in blocking solution for 1h at RT and 2 days at 4°C. After washing sections were incubated in a secondary Biotin-conjugated antibody (goat anti mouse Biotin-SP, 1:1000, #115-035-003, Dianova GmbH, Hamburg, Germany) in blocking solution for 1h at RT and 2 days at 4°C, before adding the ABC reaction reagent (Vectastain ABC Kit #PK-6100, Vector Laboratories Inc., Burlingame, CA) for 1h at RT and 1 day at 4°C. After 1 day, sections were washed before adding the DAB solution (DAB peroxidase substrate Kit #SK-4100, Vector Laboratories Inc., Burlingame, CA) for 1min. DAB reaction was stopped with purified water (ddH2O) and sections were dehydrated in the following sequence: 2 min 70 % ethanol (EtOH), 2 min 80 % EtOH, 2 min 95 % EtOH, 2 min 99,9% EtOH. Sections were cleared in Rotihistol (#6640.4, Carl Roth GmbH, Karlsruhe, Germany) until mounting in Entellan (#1.07961.0100, Merck KGaA, Darmstadt, Germany).

## Acknowledgements

Funding sources: This work was supported by National Institutes of Health grants to F.K.H. from The Eunice Kennedy Shriver National Institute of Child Health and Human Development: R01HD053889 and R01HD061575, The National Center for Research Resources: R24RR03232601; and The Office of the Director: R24OD011108. Neurological analyses on Pclo mutant rats were conducted by The Neuro-Models Facility (EJP, LBG) at UT Southwestern Medical Center, and supported by the Haggerty Center for Brain Injury and Repair. Z. Iv. was supported by grants from the Bundesministerium für Bildung und Forschung (NGFN-2, NGFNplus - ENGINE). Z. Iz. is supported by European Research Council, ERC Advanced [ERC-2011-AdG 294742]. CG is supported by the German Center for Neurodegenerative Diseases (DZNE) and DFG-SFB958. We thank Christine Römer and Ruth Ann Word for their critical comments.

## Author Contributions

Designed research: MS, EJP, LBG, JMS, JAR, CCG, ZIz, FKH; Performed research: GAM, MS, EJP, LBG, KMC, JC, PJ, HMP, AP, JMS, CB, FA, FKH; Contributed unpublished reagents/analytic tools: ZIv, ZIz, FKH, Analyzed data GAM, MS, EJP, LBG, KMC, JMS, JAR, XX, ZIv, CB, FA, CCG, ZIz, FKH; Wrote the paper GAM, EJP, KMC, MS, JMS, JAR, ZIv, CCG, ZIz, FKH.

## Supporting Information

**S1 Fig.**
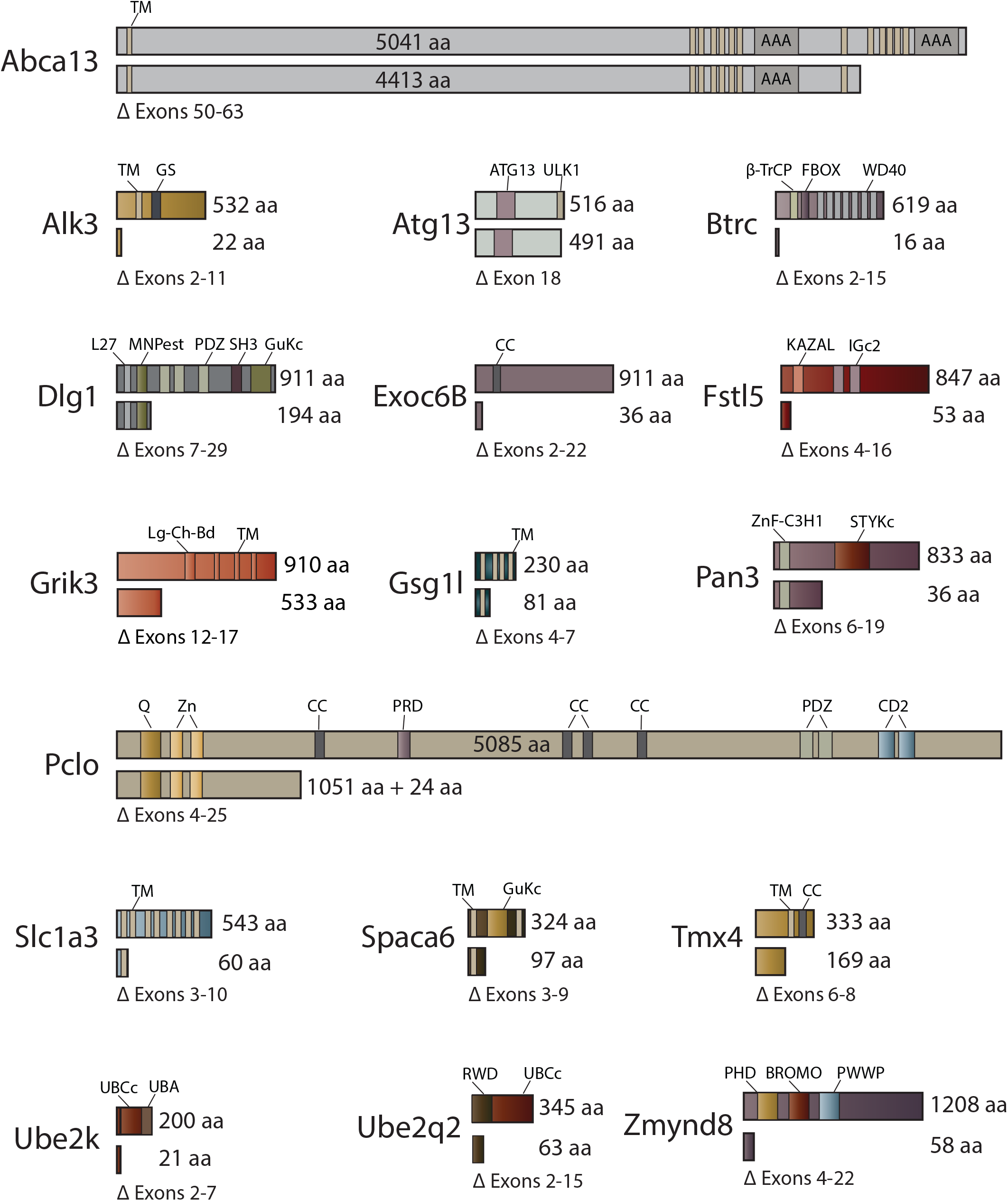
Predicted proteins produced in *Sleeping Beauty β-geo* genetrap rat strains. Exon sequences predicted to be excluded (Δ) from mRNAs encoding truncated polypeptides (aa) generated by imposed splicing to the genetrap transposon are shown below respective wildtype proteins for 17 of the 18 mutant rat strains screened for effects on reproduction. Transposon insertion within intron 2 of *Rgs22* is not predicted to truncate the Rgs22 open reading frame due to its intronic genetrap cassette inserting in the 3’ to 5’ orientation. See: S1 Table for full amino acid sequences of the 17 predicted truncated proteins encoded by respective trapped genes, which contain additional epitopes of either 3, 24 or 1319 (β-GEO) amino acids derived from the genetrap construct. TM, Transmembrane domain; AAA, ATPase Associated with a variety of cellular activities; GS, GS Motif; L27, domain in receptor targeting proteins Lin-2 and Lin-7; MN-PEST, Polyubiquitination (PEST) N-terminal domain of MAGUK; PDZ, Domain present in PSD-95; β-TrCP, D domain of beta-TrCP; FBOX, A Receptor for Ubiquitination Targets; Dlg, and ZO-1/2; SH3, Src homology 3 domain; GuKc, Guanylate kinase homologue; CC, coil coil region; KAZAL, Kazal type serine protease inhibitors; IGc2, Immunoglobulin C-2 Type; Lg-Ch-Bd, Ligated ion channel L-glutamate- and glycine-binding site; ZnF_C3H1, Zinc Finger Domain; STYKc, Protein kinase; unclassified specificity; C2, Protein kinase C conserved region 2 (CalB); UBCc, Ubiquitin-conjugating enzyme E2, catalytic domain homologue; UBA, Ubiquitin associated domain; RWD, domain in RING finger and WD repeat containing proteins and DEXDc-like helicases subfamily related to the UBCc domain; PHD, PHD zinc finger; BROMO, bromo domain; PWWP, domain with conserved PWWP motif.

**S2 Fig.**
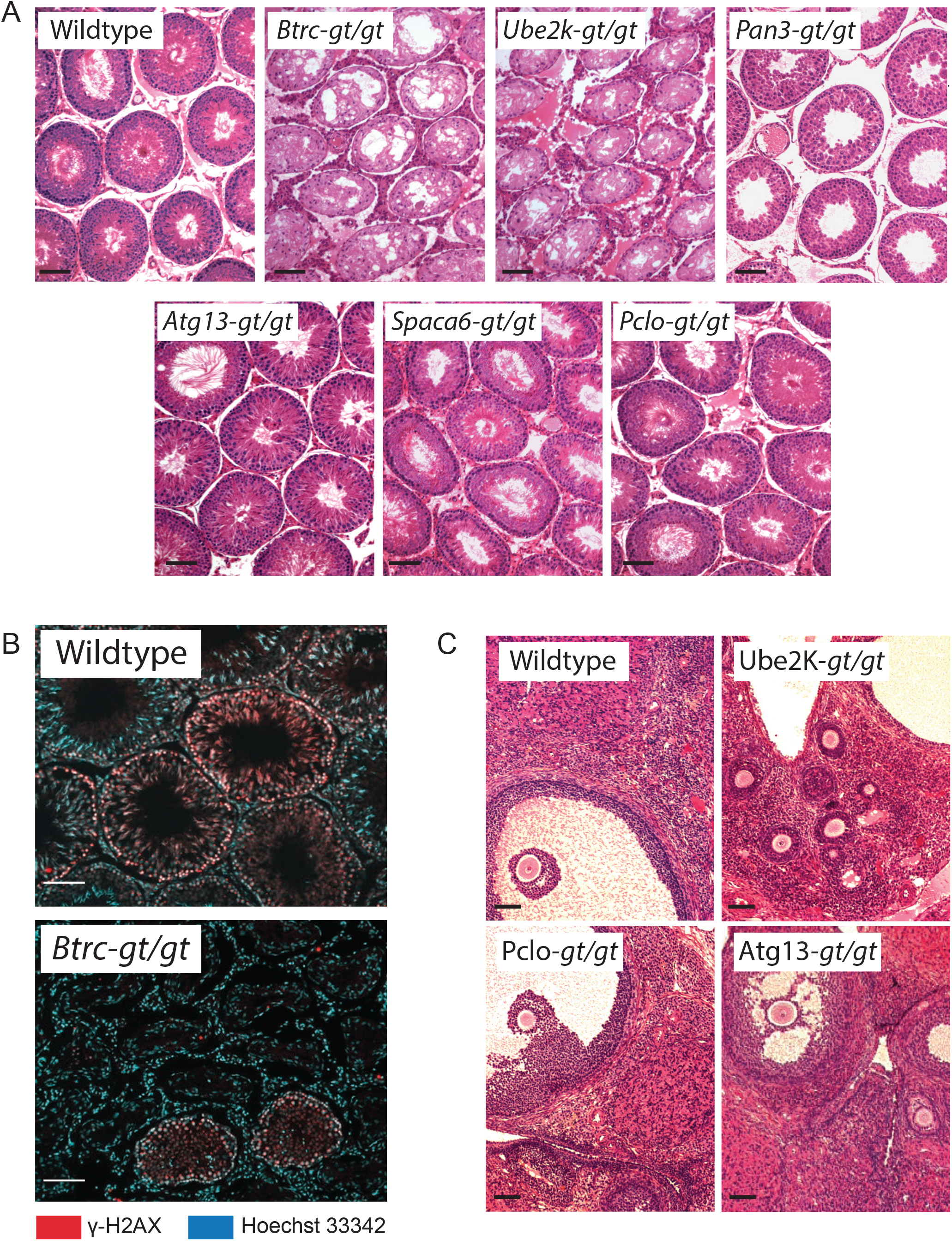
Gametogenesis defects in rats with gene-trap mutations. (**A**) H&E stained testis sections from wildtype and respective homozygous genetrap mutant rats. Scale bar, 100 μm. (**B**) Immunofluorescence labeling of cells in wildtype and mutant *Btrc*^gt/gt^ rat testis sections using an antibody to γH2AX and Hoechst 43332 dye as nuclear markers. Scale bar, 100 μm. (**C**) H&E stained ovarian sections from wildtype and respective homozygous genetrap mutant rats. Scale bar, 100 μm.

**S3 Fig.**
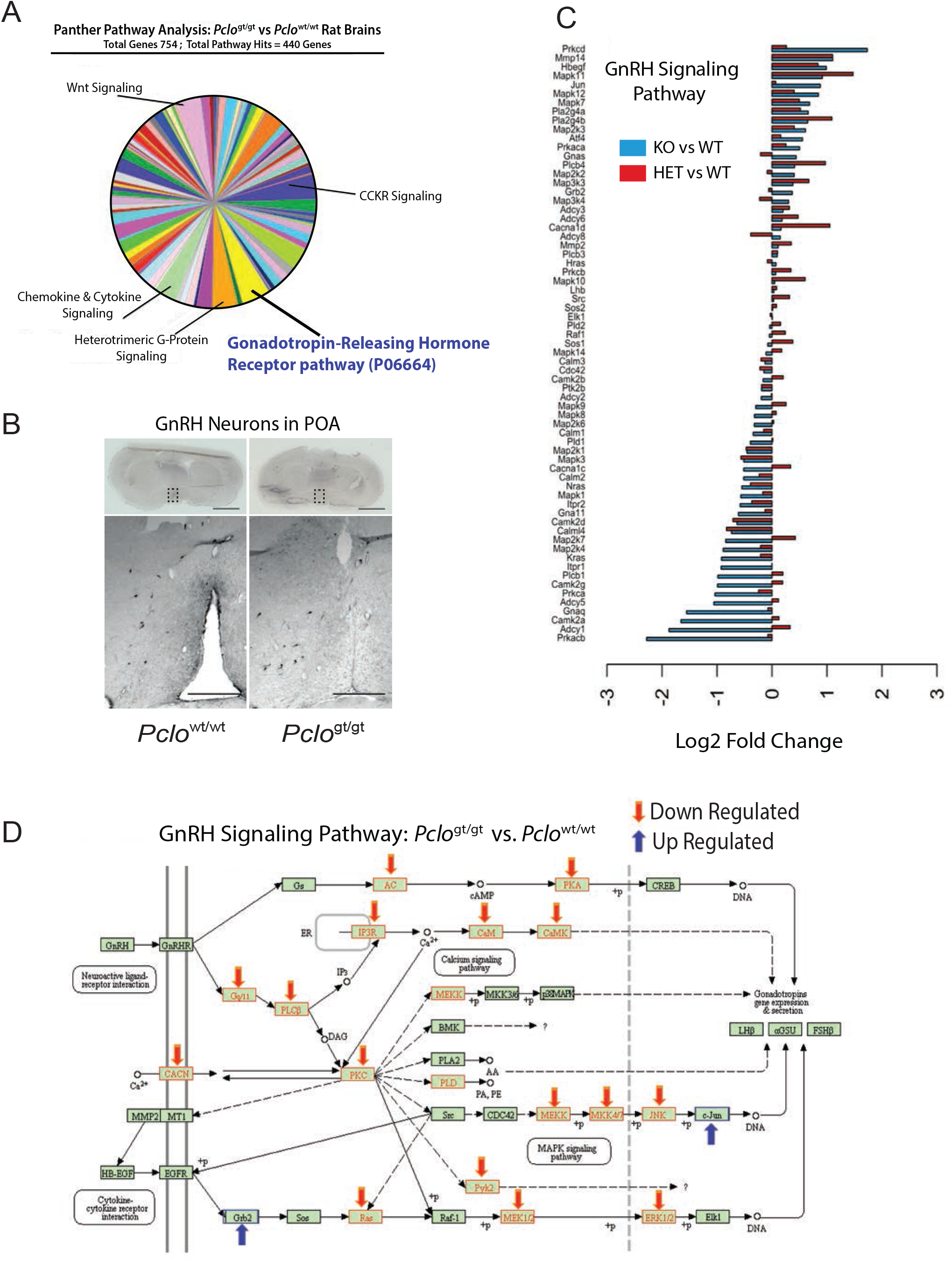
Gonadotropin Releasing Hormone (GnRH) signaling pathway genes are downregulated in the brain of mutant Piccolo rats. (**A**) Panther Pathway finder analysis on down regulated Hormonal Secretion GO: gene sets identify GnRH signaling pathway as the prominent cluster of DEGs in *Pclo*^gt/gt^ vs *Pclo*^wt/wt^ rat brains. (**B**) Coronal brain sections from Pclo^wt/wt^ or Pclo^gt/gt^ postnatal day 100 rat brains immuno-stained with GnRH antibodies. Lower panel, 5x magnification of the preoptic area (POA) located in the boxed area reveal presence of somata and GnRH positive neuron processes flanking the third ventricle. Scale bars = upper panel 0.3 cm, lower panel 1000 μm. (**C**) Relative abundance of GnRH signaling pathway GO: gene set in *Pclo*^gt/gt^ (KO) and *Pclo*^wt/gt^ (HET) vs *Pclo*^wt/wt^ (WT) rat brains. (**D**) KEGG pathway analysis predicts downregulated GnRH signaling pathways in *Pclo*^gt/gt^ rat brains vs *Pclo*^wt/wt^ rat brains. *<u>Note</u>*: See Panel C for official GnRH signaling pathway gene symbols; KEGG pathway analysis illustrates common gene acronyms.

**S4 Fig.**
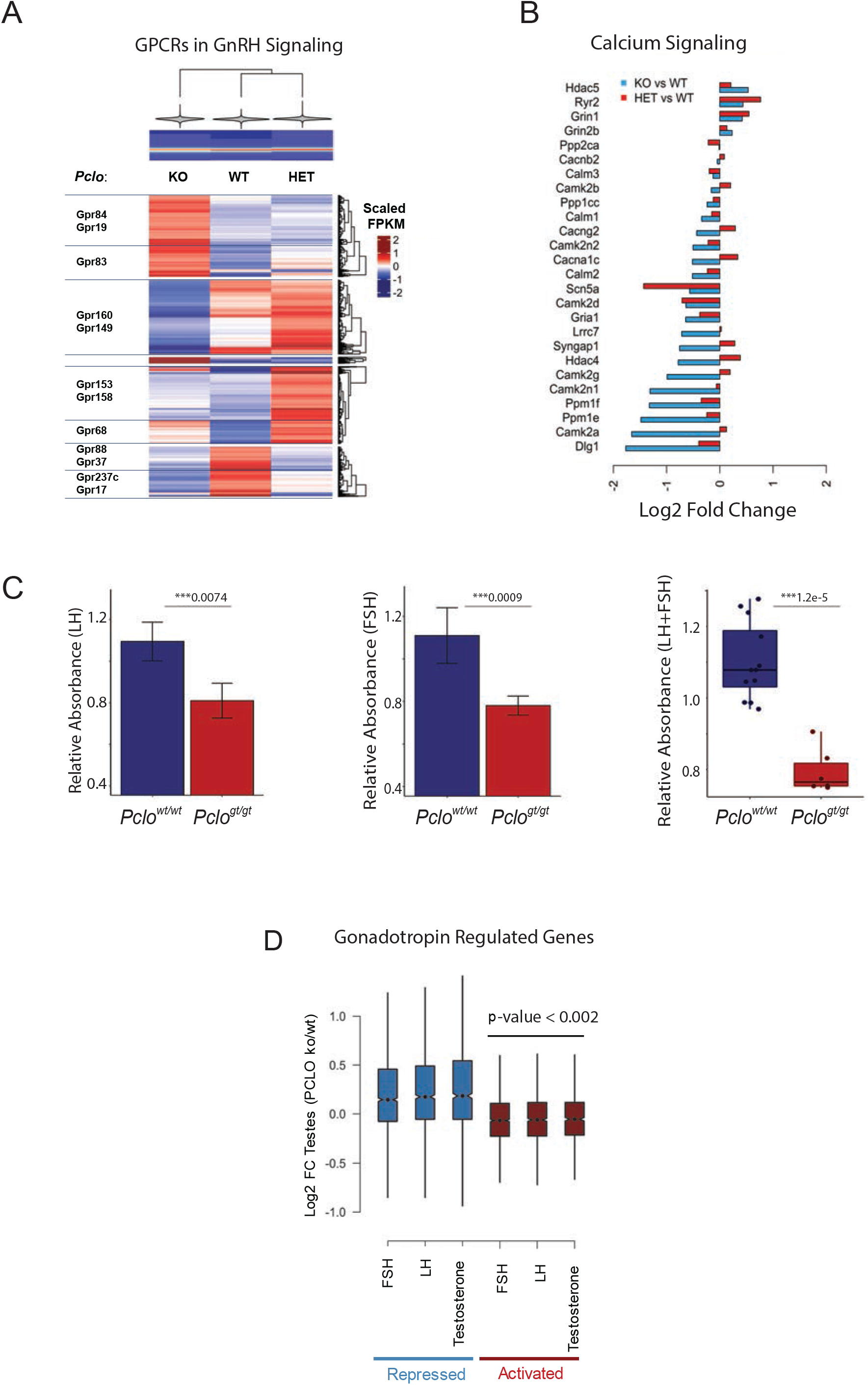
Misregulated gene networks downstream of GnRH signaling. (**A**) Relative abundance of DEGs encoding G-Protein Coupled Receptors (GPCRs) involved in conducting GnRH signaling on gonadotropes GO: gene set components in Pclo^gt/gt^ (KO) and Pclo^wt/gt^ (Het) vs Pclo^wt/wt^ (WT) rat brains. (**B**) Relative abundance of Calcium Signaling GO: gene set components in Pclo KO and HET vs WT rat brains. *Note, Dlg^wt/gt^ mutant rats displayed altered behavior and reduced fecundity (see S4 Table). (**C**) Bar plots demonstrate relative FSH (left) and LH (center) concentrations in *Pclo*^wt/wt^ and *Pclo*^gt/gt^ rats determined by ELISA at two dilutions (5x and 10x) and three technical replicates. ELISA mean ±SD absorbance values obtained from three animals (two *Pclo*^wt/wt^ and one *Pclo*^gt/gt^). Asterisks represent p-values obtained by t-test. Boxplot with jitters (right) is an alternative representation of LH and FSH levels analyzed together between *Pclo*^wt/wt^ vs *Pclo*^gt/gt^ rats. Asterisks represent p-values obtained by Wilcox test. Each dot represents a replicate (biological/technical). (**D**) Relative abundance of Gonadotropin Regulated Genes in Pclo KO vs WT rat testes (PCLO ko/wt) that are Repressed or Activated by FSH, Follicle Stimulating Hormone; LH, Luteinizing Hormone, T, Testosterone (Zhou et al., 2010).

**S1 Table. Gene Trap Mutations in Rat Strains Screened for Reproduction Phenotypes**

**S2 Table. Reproduction Phenotypes in Rats with Sleeping Beauty Gene Trap Mutations**

**S3 Table. Body, Testis and Epididymal Weights in Sleeping Beauty Mutant Rats**

**S4 Table. Mutant Rat Phenotypes in Current Study* Compared Across Species.**

**S5 Table. RNA Sequencing: Brain and Testes Transcriptomes *Pclo* Mutant Rats**

**S6 Table. Gene Ontology: Brain and Testes Transcriptomes *Pclo* Mutant Rats**

**S1 Video. Female WT & Male WT Rats**

**S2 Video. Female *Pclo* KO & Male WT Rats**

**S3 Video. Male *Pclo* KO & Male WT Rats**

**S4 Video. Male WT & Male WT Rats**

**S5 Video. Male WT & Male *Pclo* HET Rats**

**S6 Video. Male *Pclo* KO & Male *Pclo* WT Rats**

